# *Verticillium dahliae* effector VDAL protects MYB6 from degradation by interacting with PUB25/26 E3 ligases for enhancing Verticillium wilt resistance in *Arabidopsis*

**DOI:** 10.1101/2021.03.26.437179

**Authors:** Aifang Ma, Dingpeng Zhang, Guangxing Wang, Kai Wang, Zhen Li, Yuanhui Gao, Hengchang Li, Chao Bian, Jinkui Cheng, Yinan Han, Shuhua Yang, Zhizhong Gong, Junsheng Qi

## Abstract

Verticillium wilt is a severe plant disease that causes massive losses in multiple crops. Increasing the plant resistance to Verticillium wilt is a critical challenge worldwide. Here, we report that the hemibiotrophic *Verticillium dahliae* (*V. dahliae*)-secreted Asp f2-like protein VDAL causes leaf wilting when applied to cotton leaves *in vitro,* but enhances the resistance to *V. dahliae* when overexpressed in *Arabidopsis* or cotton without affecting the plant growth and development. VDAL protein interacts with *Arabidopsis* E3 ligases PUB25 and PUB26 (PUBs) and is ubiquitinated by PUBs *in vitro*. However, VDAL is not degraded by PUB25 or PUB26 *in planta*. Besides, the *pub25 pub26* double mutant shows higher resistance to *V. dahliae* than the wild type. PUBs interact with the transcription factor MYB6 in a yeast two-hybrid screen. MYB6 promotes plant resistance to Verticillium wilt while PUBs ubiquitinate MYB6 and mediate its degradation. VDAL competes with MYB6 for binding to PUBs, and the role of VDAL in increasing Verticillium wilt resistance depends on MYB6. Taken together, these results suggest that plants evolute a strategy to utilize the invaded effector protein VDAL to resist the *V. dahliae* infection without causing a hypersensitive response (HR), Alternatively, in order to take nutrients from host cells, hemibiotrophic pathogens may use some effectors to keep plant cells alive during its infection in order to take nutrients from host cells. This study provides the molecular mechanism for plants increasing disease resistance when overexpressing some effector proteins without inducing HR, and may promote searching for more genes from pathogenic fungi or bacteria to engineer plant disease resistance.

**One-sentence summary:** Ectopically expressed VDAL in *Arabidopsis* alleviates the degradation of a positive disease response factor MYB6 through its interaction with PUB25/26 E3 ligases.

## INTRODUCTION

The plant immune system confers effective protection against diverse pathogens. The activation of this system requires pathogen-derived molecules, which are classified into two categories (Jones and Dangl, 2006; Eckardt, 2017; Wang et al., 2020b; Zhou and Zhang, 2020). The conserved pathogen-associated molecular patterns (PAMPs) are molecules belong to the first category, which are perceived by specific plant pattern-recognition receptors (PRRs) at the cell surface (Kunze et al., 2004; Zipfel et al., 2006; Sun et al., 2013; Zhou and Zhang, 2020). This PRR-initiated immunity, also known as PAMP-triggered immunity (PTI), generally occurs upon the initial contact of the pathogen with the plant cell, thus constituting the first layer of the immune response (Afzal et al., 2011; Singh et al., 2014; Bigeard et al., 2015; Wang et al., 2020b; Zhou and Zhang, 2020). PTI occurs quickly upon the perception of PAMPs when pathogens infect plants, and causes a series of rapid immune responses, such as a reactive oxygen species (ROS) burst, the activation of mitogen-activated protein kinase (MAPK) activity, callose accumulation, the induced expression of disease resistance genes, and changes in hormone levels (Boller and Felix, 2009; Boller and He, 2009; Qi et al., 2018; Wang et al., 2020b). The second category is called as pathogen secreted effectors that can be delivered into plant cells by various mechanisms (Liu et al., 2013; Rajamuthiah and Mylonakis, 2014; Stotz et al., 2014; Cui et al., 2015; Betsuyaku et al., 2018). Although some effectors reprogram the transcriptional profile in the host cells to create the niche permissive for infection (Boch et al., 2009; Boch and Bonas, 2010; Christian et al., 2010), most effectors suppress PTI by directly engaging specific host proteins via physical interactions or biochemical modifications with unique enzymatic activity (Block et al., 2014; Lin et al., 2016). To resist invasion of pathogen effectors, plants employ another defense response known as effector-triggered immunity (ETI), often accompanied by a hypersensitive response (HR), at the infection site, to limit pathogen infection (Jones and Dangl, 2006; Dodds and Rathjen, 2010; Wang et al., 2020b; Zhou and Zhang, 2020). ETI and PTI can reinforce each other to increase the plant resistance to pathogens (Ngou et al., 2021; Yuan et al., 2021b; Yuan et al., 2021a).

*Verticillium dahliae* (*V. dahliae*) is a soil-borne hemibiotrophic fungal pathogen that causes Verticillium wilt in more than 200 host species, including important economic crops (Klosterman et al., 2009; Njoroge et al., 2009; Wheeler and Johnson, 2016). Verticillium wilt causes significant economic losses to agriculture worldwide. For instance, cotton (*Gossypium hirsutum*) production in many areas of the world is severely threatened by this disease (Zhao et al., 2014; Han et al., 2019). As *Arabidopsis thaliana* can also be infected by this fungal pathogen (Veronese et al., 2003; Qin et al., 2018), it is often used as a model plant to study the molecular mechanism underlying Verticillium wilt resistance.

*V. dahlia*e infection was characterized by the association of fungal hyphae with the plant’s woody vascular tissues, where fungicides cannot readily reach, thus may account for the difficulty in controlling Verticillium wilt (Vallad and Subbarao, 2008; Zhao et al., 2014; Deng et al., 2015). As one of the fungal pathogen weapons, secreted proteins are thought to play critical roles in the pathogenicity of *V. dahliae* (Klosterman et al., 2011; Lo Presti et al., 2015). These secreted proteins usually act as effectors that are employed by pathogens to overcome plant defense. Exposure to these proteins often leads to PTI or ETI in the host cells (Dodds et al., 2004; Doehlemann et al., 2009; Lo Presti et al., 2015). *V. dahliae* secretes more than 700 putative effectors, including more than 100 small cysteine-rich potential effectors (Klosterman et al., 2009; Klosterman et al., 2011), but only a few were functionally studied. For example, the tomato cell-surface-localized immune receptor Ve1 is activated by the *V. dahliae*-secreted protein AVe1 (Avirulence on Ve1) to trigger plant immune responses (Fradin et al., 2009; Deng et al., 2015). Effector isochorismatase (VdIsc1) hydrolyses isochorismate [the direct precursor of salicylate (SA)] to suppress SA-mediated defense (Liu et al., 2014b), while the effector VdSCP41 (*V. dahliae* secretory protein 41) interacts with plant-specific transcription factors CBP60g (CALMODULIN BINDING PROTEIN 60) and SARD1 (SYSTEMIC ACQUIRED RESISTANCE DEFICIENT 1) further inhibits transcriptional activity of CBP60g (Qin et al., 2018). Another study found that infiltration of *V. dahliae* secreted protein PevD1 to cotyledons enhances cotton resistance and the defense response to the *V. dahliae* infection (Bu et al., 2014).

Protein ubiquitination, a post-translational modification, is required for many signaling pathways involved in numerous essential cellular processes including plant growth, development and stress response in eukaryotic cells (Kong et al., 2015; You et al., 2016; Yang et al., 2017; Ye et al., 2018; Zhou. et al., 2018). Protein ubiquitination reaction is catalyzed by three enzymes, the ubiquitin (Ub)-activating E1, the Ub-conjugating E2, and the Ub-ligase E3 (Yu et al., 2016), and this modification is also critical for defense responses in plants (Dombrecht et al., 2007; Tong et al., 2017; Furniss et al., 2018). E3 ligases that dictate the specificity of the substrate for ubiquitination are highly diverse (Kosarev. et al., 2002), and can be classified into 4 groups, including HECT (Homologous to E6-associated protein C-Terminus), RING (Really Interesting New Gene), U-box and Cullin (Cul)-RING ligase (CRLs) (Stone, 2014; Liao et al., 2017). Plant U-box (PUB) E3 ligases in *Arabidopsis* have 64 members (Mudgil et al., 2004). All PUBs contain a high conserved U-box domain, which is required for interaction with E2 protein (Pringa et al., 2001) and essential for PUB ligase activity (Ohi et al., 2003; Zeng et al., 2004). Only a limited number of PUBs have been functionally studied in plants. For example, PUB12/13 participate ABA signal pathway by degrading the key negative ABA coreceptor ABA INSESSITIVE 1 (ABI1) (Kong et al., 2015). PUB10 and PUB2/4 plays roles in jasmonic acid and cytokinin response (Jung et al., 2015; Wang et al., 2017b). Since PUB12 and PUB13 were also found to target FLAGELLIN-SENSITIVE 2 (FLS2) for its degradation (Lu. et al., 2011), more and more lines of evidence indicate that PUBs play important roles in plant disease resistance. PUB22, PUB23 and PUB24 appears to negatively regulate plant PTI, and the mutants defective in these three genes exhibit higher PAMP-induced oxidative burst and inhibit bacterial growth (Stegmann et al., 2012). A recent study indicates that PUB25/26 target and degrade non-activated BOTRYTIS-INDUCED KINASE 1 (BIK1) to negatively regulate immunity in *Arabidopsis* (Wang et al., 2018a). These two E3 ligases also promote *Arabidopsis* freezing tolerance by degrading the cold signaling negative regulator MYB15 (Wang et al., 2019). PUB25/26 negatively regulate petal growth in a spatial- and temporal-specific manner (Li et al., 2021). GhPUB17 from cotton plays a negative role in plant resistance to *V. dahliae*, and its E3 ligase activity can be inhibited by a cyclophilin homolog GhCyP3 (Qin et al., 2019). In other plant species, PUB ligases were found also to be linked to biotic stress (Ishikawa et al., 2014; Jiao et al., 2017).

Transcription factors (TFs) are important for gene expression. According to different DNA binding domains, they are divided into different families, such as MYC, MYB, bZIP and bHLH family (Pabo and Sauer, 1992). The MYB TFs constitute the largest TF family among all eukaryotic organisms (Riechmann et al., 2000). The N-terminus of MYB TF contains a repetitive sequence consisting of MYB domain with about 52 amino acids that can form three α helix structures. The second and third α helix of each MYB domain can form helix-turn-helix (HTH), which can bind to the DNA trench (Ogata et al., 1994). According to the number of MYB domains, MYB TFs can be divided into four categories, called 1R (R1/2, R3-MYB), 2R (R2R3-MYB), 3R (R1R2R3-MYB) and 4R (Dubos et al., 2010). Most of MYBs in plants belong to 2R (R2R3-MYB). There are 138 R2R3-MYBs identified in *Arabidopsis* (Katiyar et al., 2012). Acting as transcription activators or inhibitors, MYB TFs participate in plant primary and secondary metabolism, cell fate and properties, plant growth and development, and plant adaptation to different stresses (Preston et al., 2004; Zhong et al., 2007; McCarthy et al., 2009; Dubos et al., 2010; Liu et al., 2015). In flg22-treated *Arabidopsis* plants, the expression of *MYB12*, a positive regulator in flavanone synthesis, was up-regulated, while the expression of *MYB4* was down-regulated, suggesting an antagonistic effect between two MYBs to coordinate plant immune response (Schenke et al., 2011; Jin et al., 2014). MYB96 modulates plant immune response by regulating salicylic acid (SA) biosynthesis (Seo and Park, 2010). MYB30 is involved in regulating plant hypersensitive response (Marino et al., 2013). In an early study, the DNA binding motif of MYB6 or MYB7 in *Arabidopsis* was determined (Roger W.parish and Li, 1995). The expression of *MYB6* was induced by JA or SA treatment in *Arabidopsis* (Yanhui et al., 2006). However, whether MYB6 is involved in regulating the immunity of *Arabidopsis* to *V. dahliae* or plays other biological roles has not been reported yet.

To test the function of *V. dahliae* secreted proteins, we generated transgenic *Arabidopsis* and cotton plants ectopically expressing the *V. dahliae-*secreted Asp f2-like protein (VDAL). These transgenic plants exhibited significant resistance to this fungal pathogen infection, without any growth or developmental defects. Previously, several effectors such as PsCRN161 from *Phytophthora sojae*, SSB (Xoc), and Harpin (xoo) from *Xanthomonas oryzae pv. oryzicola* were reported to increase plant resistance to pathogens without retarding plant growth when overexpressed in plants, however, the molecular mechanisms are largely unknown (Peng et al., 2004; Rajput et al., 2015; Cao et al., 2018). In this study, we show that VDAL interacts with the E3 ligases PUB25 and PUB26, but was not degraded in *Arabidopsis*, while PUB25 and PUB26 target a positive regulator of plant disease resistance MYB6 for its degradation. Our results suggest that VDAL competes with MYB6 to bind PUB25 and PUB26, and increases the accumulation of MYB6, thus improving disease resistance in *Arabidopsis*. These results suggest that plants not only use ETI and PTI to produce ROS (Wang et al., 2020b; Zhou and Zhang, 2020), but also evolute a strategy to utilize some effectors such as VDAL without the sacrifice of cell death to resist the *V. dahliae* infection, or pathogens take advantage of the effectors not killing host cells immediately in order to fully exploit the nutrients of host cells. *VDAL* may represent an ideal candidate gene for improving Verticillium wilt resistance in crops without affecting plant growth or yields.

## RESULTS

### Identification of VDAL, a protein secreted by *V. dahliae*

In an effort to identify proteins secreted by *V. dahliae* that might be responsible for Verticillium wilt symptoms in plants, we identified a protein named VDAL (*Verticillium dahliae* secreted Aspf2-like protein) from the *V. dahliae* strain Vd991, which is homologous to human Pra1 and Aspf2 (Supplemental Figure 1A). Aspf2 is present in many fungi, including animal fungal pathogens such as *Aspergillus fumigatus* (Supplemental Figure 1A). Sequence analysis revealed that VDAL contains the putative Zn^2+^-binding motif HARxH, which is present in many metalloproteases, and a secretion signal peptide (SP) in its first 23 residues (Supplemental Figure 1B). VDAL is very conserved in some fungi such as *Fusarium*, *Colletotrichum*, *Metarhizium*, and *Purpureocillium* (Supplemental Figure 2A and Supplemental data 1), which may have a general role as an effector. We performed an invertase secretion assay to investigate whether this secretion signal peptide is functional. Specifically, we fused the putative VDAL signal peptide with the sucrose invertase sequence in pSUC2-MSP and introduced the resulting construct into yeast strain YTK12, which is defective in invertase activity (SUC2), to generate the pSUC2-VDAL-SP fusion protein (Supplemental Figure 1C). As expected, this strain and a control strain harboring the empty vector (pSUC2^-MSP^) both grew robustly on CMD-W Trp dropout medium containing 0.1% glucose and 2% sucrose (Figure 1A). However, only strain YTK12-VDAL-SP gained the ability to grow on YPRAA medium with raffinose as the sole carbon source, indicating that invertase was secreted by this strain (Figure 1A). In agreement with this result, strain YTK12-VDAL-SP reduced 2, 3, 5-Triphenyltetrazolium chloride in the medium into red formazan (Figure 1A), further confirming that invertase is secreted by VDAL SP. To further verify the secreting activity of VDAL SP, we deleted and mutated the critical amino acids of VDAL SP, named SP-mut-1 to SP-mut-5, respectively (Supplemental Figure 1D), and performed the invertase secretion assay to investigate whether theses mutated SPs are functional. The N-terminus of either *Phytophthora sojae* Avr1b or *Magnaporthe oryzae* Mg87 was used as a positive or negative control, respectively (Fang et al., 2016). The results showed that except for the SP-mut-2 that is a little weaker than the wild type, the secreting activity of other different mutated SPs was greatly compromised, suggesting that SP has secreting ability (Supplemental Figure 1E). Since VDAL is an M35-like metalloprotease that might have zinc-binding activity, we tested its zinc-binding activity. Like Pra1, VDAL could bind zinc *in vitro* (Supplemental Figure 1F and 1G). To determine whether VDAL causes leaf wilting, we dipped cotton leaf petioles into a solution of VDAL-HIS recombinant protein that was expressed and purified from *Escherichia coli* (*E. coli*). Due to the failure of purifying the full-length VDAL protein in *E. coli,* the SP was removed from VDAL protein in this study. Incubation with 3 ppm or 5 ppm of the VDAL-HIS solution for 12 h led to obvious leaf wilting, while the same treatment of mock treatment (buffer that dissolve VDAL-HIS protein containing) did not cause any visible wilting. We recorded water loss after 12 h of incubation and photographed the leaves after 24 h of incubation, the quantification indicated the water loss in VDAL-HIS treatment was significantly higher than the mock treatment (Figures 1B and 1C). The wilted leaves failed to recover after being transferred to water, suggesting that cell death occurred in these leaves due to the VDAL treatment.

**Figure 1.**
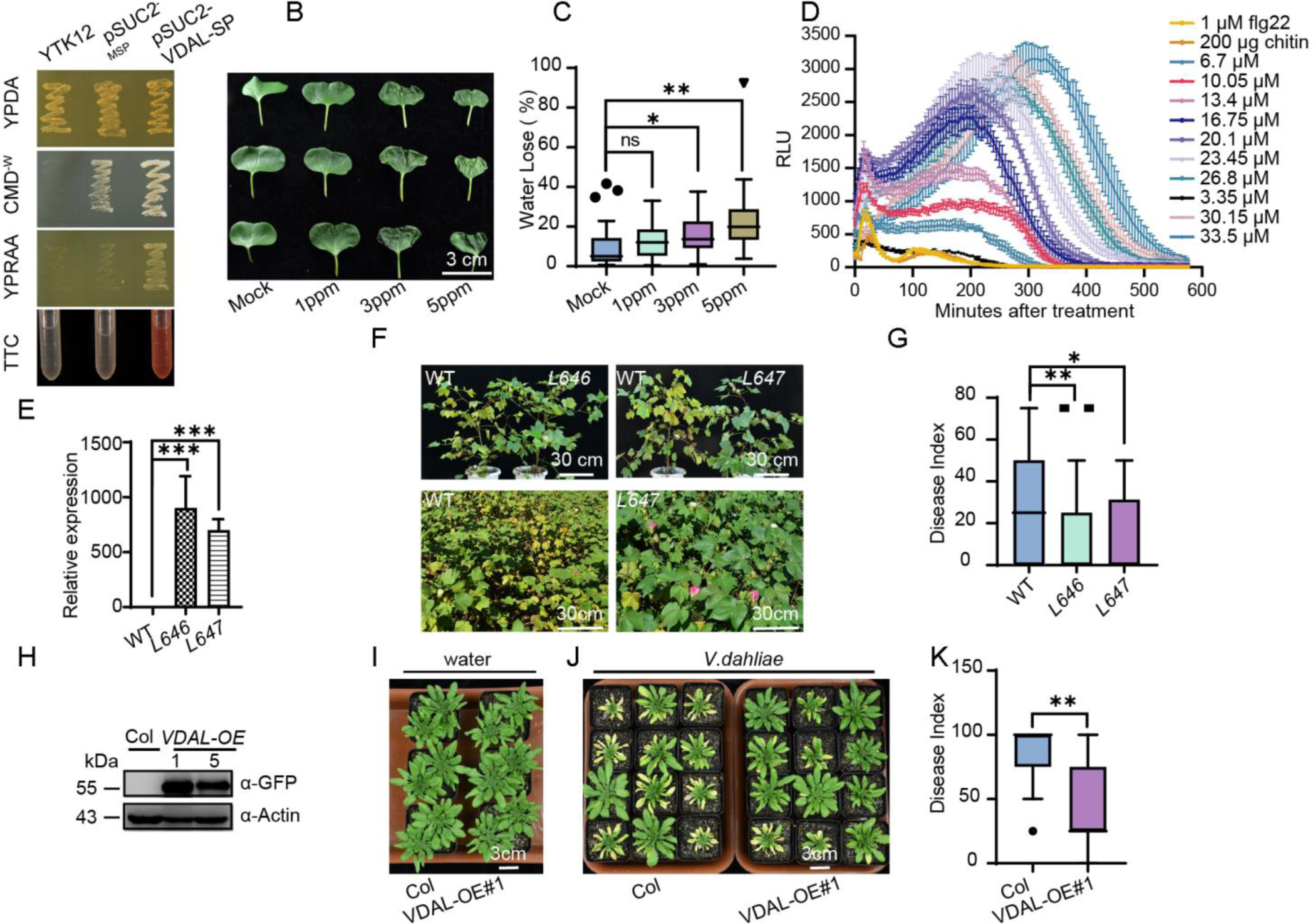
Plants overexpressing *VDAL* are resistant to *Verticillium* wilt caused by *V.dahliae*. **(A)** The signal peptide (SP) of VDAL is functional. CMD^-W^ medium for positive clone screening;YPRAA medium for verification of secretion activity on invertase; TTC assay for the test of secreted invertase activity. Note that only the strain expressing the VDAL SP fusion gained the ability to catabolize raffinose, and reduce 2, 3, 5-triphenyltetrazolium chloride (TTC) into red formazan. Both YPRAA and TTC indicate the successful secretion of the invertase. **(B)** VDAL protein causes leaf wilt. Cotton leaves became wilt after the petiole was soaked in different concentration of VDAL protein solution for 24 h, and then photos were taken. **(C)** The water loss during leaf wilting. Leaves’ weight was measured at the beginning and at 12 h after post treatment. **(D)** Luminol-based assay show that VDAL can elicit ROS burst with increasing concentration of VDAL protein. Flg22 and chitin were used as positive controls. **(E)** The expression level of *VDAL* in two corresponding transgenic cotton lines. **(F-G)** Cotton lines exhibited tolerance to Verticillium wilt caused by *V. dahliae* in the field. **(F)** Upper: two transgenic lines (*L646* and *L647*) and wild type plant were removed from the same field infected with *V. dahliae*, and replanted in the pots for taking photos; Down: photos for the wild type cotton plants and transgenic *L647* plants growing in the same field infected with *V. dahliae*. **(G)** Statistical analysis of the disease index in the above field condition, count with at least 25 plants. **(H)** Immunoblot analysis of VDAL in two transgenic *Arabidopsis thaliana* lines overexpressing *VDAL.* Total proteins of 10-day-old seedlings were extracted and detected with anti-GFP antibodies. Actin was an equal loading control. **(I-K)** VDAL transgenic *Arabidopsis* plants show resistance to Verticillium wilt caused by *V. dahliae*. The plants grown in greenhouse for two weeks were dipped into the *V. dahliae* spore suspension or water for 5 minutes. Photos were taken at 20-day post infection (20 dpi). **(K)** Statistical analysis of the disease index of *VDAL-GFP* and Col, count with at least 15 plants. *, **and *** in **(C), (E), (G)** and **(K)** represent significant difference (P<0.05), highly significant difference (P<0.01) and extremely significant difference (P<0.001), respectively, in Student’s t-test. The experiments were repeated independently three times with similar results.

We reasoned that the VDAL-incubated leaves might have exhibited a HR to VDAL and overproduced ROS. We therefore measured ROS production in VDAL-treated *Arabidopsis* leaves. Bacterial flagellin and fungal chitin are well-known PAMPs that induce extracellular ROS bursts (Wang et al., 2020b). We detected ROS production in *Arabidopsis* leaves treated with VDAL using a luminol-based assay, with the bacterial flagellin flg22 and chitin used as positive controls. As shown in Figure 1D and Supplemental Figure 1H, similar to flg22 and chitin, VDAL elicited a ROS burst, with higher levels of ROS produced as its concentration increased. Combined together, these results indicate VDAL is a pathogenic elicitor protein that can cause cotton leaf wilting and ROS burst.

### Plants expressing VDAL are resistant to infection by *V. dahliae*

The presence of a functionally secreted SP in VDAL suggests that this protein is secreted into the extracellular space or even into host cells by *V. dahliae*. Some secreted proteins such as the bacterial harpins (Peng et al., 2004), Verticillium Ave1 (Castroverde et al., 2016; Castroverde et al., 2017), SSB protein from *Xanthomonas oryzae* pv. *oryzicola* (Xoc, SSB_Xoc_) (Cao et al., 2018) and the Crinkler (CRN) effector PsCRN161 from *Phytophthora sojae* (Rajput et al., 2015) have been used to activate plant immunity by stably expressing them in transgenic plants without inducing a HR. Except for PsCRN161 that can suppress cell death induced by other elicitors (Rajput et al., 2015), all other abovementioned effectors can induce HR like VDAL when applied on plants. We generated transgenic cotton (*Gossypium hirsutum*) lines *L646* and *L647* stably expressing VDAL (Figure 1E) and examined their tolerance to wilt disease in the field. The most transgenic plants did not show any growth or developmental phenotypes compared to non-transgenic wild-type plants in the normal field. However, the transgenic plants exhibited increased tolerance to Verticillium wilt, with lower disease index than the wild type under field conditions (Figures 1F and 1G). We also test more different VDAL transgenic cotton lines and obtained the similar results (Supplemental Figure 1I).

We then generated transgenic *Arabidopsis* plants to examine the potential role of VDAL in triggering plant immune responses. Of the various transgenic *Arabidopsis* lines generated, we selected two lines (#1 and #5) for further study. VDAL protein was readily detected in both transgenic lines by immunoblotting (Figure 1H). Like in cotton, the expression of VDAL in *Arabidopsis* did not alter plant morphology or other traits such as growth, leaf color or the timing and rate of reproduction under normal growth conditions (Supplemental Figure 1J). We then examined the susceptibility of these plants to *V. dahliae* infection. We carefully removed 2- to 3-week-old seedlings from the soil, washed the roots to remove any attached soil, and incubated them in a 1×10^6^ conidia/ml spore suspension of *V. dahliae* strain Vd991 for 5 min. We transferred the inoculated plants to new pots filled with soil and observed the development of disease symptoms. During the first 7 days post inoculation (dpi), all plants resumed growth and no disease symptoms were observed. At 15 dpi, the leaves of inoculated wild-type plants began to become discolored, and no plants survived for more than 30 dpi (Figure 1J). By contrast, plants expressing VDAL did not show serve discernable symptoms at 15 dpi, and leaf discoloration did not appear until 20 dpi. Quantification of disease symptoms by determining the disease index (Figure 1K) of the plants at 20 dpi indicated that the VDAL transgenic lines exhibited significantly weaker disease symptoms than wild-type plants. Thus, both cotton and *Arabidopsis* plants expressing VDAL enhanced resistance to Verticillium wilt.

### VDAL interacts with the U-box ubiquitin E3 ligases PUB25 and PUB26

To determine the mechanism of VDAL-induced Verticillium wilt resistance, we examined the subcellular localization of VDAL by expressing a VDAL-GFP fusion protein in *Arabidopsis* protoplasts. Robust expression of the fusion protein was detected at 16 h after transfection. The distribution pattern of VDAL-GFP indicated that VDAL localized to the cytosol, with some dots of signal observed (Supplemental Figure 1K). A similar localization pattern was observed in *VDAL-GFP* overexpression *Arabidopsis* plants (Supplemental Figure 1L).

To identify the host proteins targeted by VDAL, we transiently expressed a VDAL-MYC fusion protein in *Arabidopsis* protoplasts for 16 h, and incubated total proteins extracted from transfected cells with agarose beads coated with MYC-specific antibodies for 2.5 h. After extensive washing, the eluted bound proteins were concentrated and analyzed by SDS-PAGE, followed by silver staining to visualize proteins co-immunoprecipitated by VDAL-MYC. Mass spectrometry analysis identified several co-immunoprecipitated proteins, including PUB26 (Supplemental Data Set 1). PUB25 and PUB26 are two closely homologous U-box proteins. Previous studies found that PUB25 and PUB26 target non-activated BIK1 during plant immunity and MYB15 during the freezing response (Wang et al., 2018a; Wang et al., 2019), thus we focused on the relationship between VDAL and these two E3 ligases.

To determine whether VDAL directly interacts with these two U-box proteins, we performed a yeast two-hybrid assay using VDAL fused to the activation domain (AD) of the yeast transcriptional activator Gal4, and either PUB25 or PUB26 fused to its DNA-binding domain (BD) (Li et al., 2020). We constructed a series of yeast strains by transforming yeast Gold cells with various combinations of plasmids, including a pair that expressed fusions of BIK1 and PUB26, two established interacting proteins (Wang et al., 2018a). As expected, cells expressing the BD-PUB26 and AD-BIK1 fusions were able to grow on the reporter medium used to measure interactions (Figure 2A). Interactions were detected between VDAL and PUB25 or PUB26 (Figure 2A). By contrast, no interaction was detected in yeast strains co-expressing the fusion proteins with two Arabidopsis proteins (AT4g20360, a plastid localized EF-Tu translation elongation factor, and AT3g44110, a co-chaperon DNAJ protein), and only the DNA-binding motif or the activation motif (Figure 2A), indicating that VDAL specifically interacts with PUB25 and PUB26.

**Figure 2.**
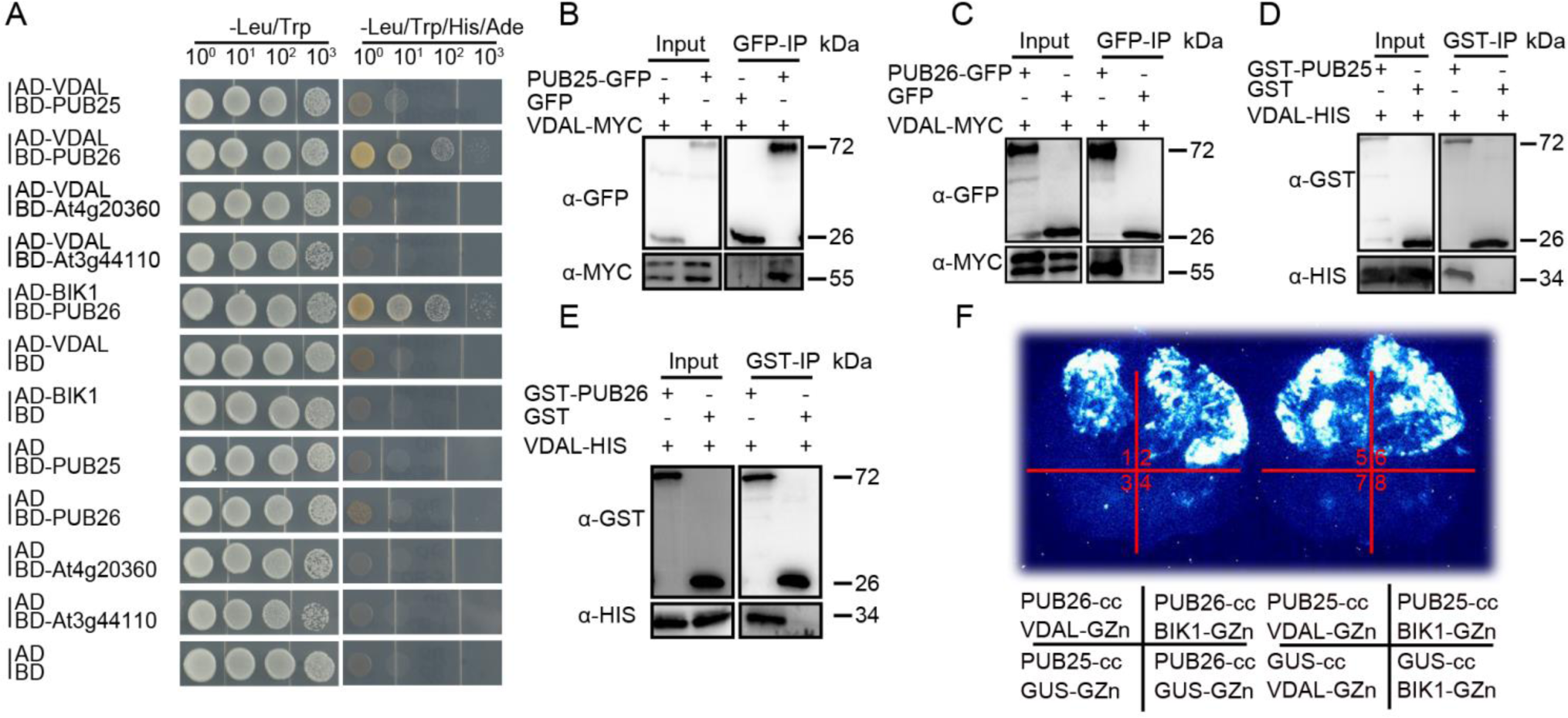
VDAL interacts with PUB25 and PUB26, two predicted plant U-box ubiquitin E3 ligases. **(A)** VDAL interacts with PUB25 and PUB26 in yeast two-hybrid (Y2H) assay. The interaction of BD-PUB26 and AD-BIK1 is used as a positive control. BD-At4g20360 and BD-At3g44110 are used as negative controls. Yeast cells were grown on different medium for four days. AD: Gal4 activation domain; BD: Gal4 DNA-binding domain. **(B)** and **(C)** VDAL interacts with PUB25 **(B)** and PUB26 **(C)** in Co-IP assay. *VDAL-MYC* and *PUB25-GFP* or *PUB26-GFP* plasmids were co-transfected into *Arabidopsis* protoplasts, and incubated for 16 h. Total proteins were extracted and immunoprecipitated with anti-GFP beads, then immunoblotting analysis was carried out with anti-MYC and anti-GFP antibodies. **(D)** and **(E)** VDAL interacts with PUB25 **(D)** and PUB26 **(E)** in the pull-down assay. Proteins purified from *E. coli* were immunoprecipitated with anti-GST beads and detected with anti-GST and anti-HIS antibodies. **(F)** VDAL interacts with PUB25 and PUB26 in the split firefly luciferase complementation imaging (LCI) assay. Different plasmids combinations were transiently expressed in tobacco leaves for 72 h, then images were collected by a CCD camera.

To identify which fragment of VDAL interacts with full-length PUB25 and PUB26 and their ARM domains, we designed two types of deletion mutants of VDAL (Supplemental Figure 3A). VDAL (1-59) interacted with full-length PUB26 but not PUB25, whereas VDAL (60-297) interacted with both full-length PUB proteins (Supplemental Figure 3B). The other deletion mutations did not interact with the full-length PUBs or their ARM domains (Supplemental Figure 3B).

We further verified the interactions between VDAL and the two ubiquitin ligases by co-immunoprecipitation (Co-IP), pull-down and firefly luciferase complementation imaging assays (LCI). We subjected total proteins from protoplasts transiently expressing VDAL-MYC and PUB25-GFP or PUB26-GFP to Co-IP assays using beads coated with anti-GFP antibodies. The protein corresponding to VDAL-MYC was readily detected in lysates co-expressing these two proteins. By contrast, these proteins were not detected in lysates from cells co-expressing GFP and VDAL-MYC (Figures 2B and 2C). These results confirm the specific interactions between VDAL and the PUBs. Similar results were obtained in a pull-down assay. We purified GST-PUB25, GST-PUB26, GST and VDAL-HIS from *E. coli*. Pull-down assays using GST beads indicated that GST-PUB25 or GST-PUB26, but not GST, was able to pull down VDAL-HIS (Figures 2D and 2E). Robust interactions were also observed in the LCI assays. Strong luminescence was detected only in leaf sections expressing both VDAL and the PUB25 and PUB26 fusion proteins (Figure 2F, Supplemental Figures 4A and 4B). Here, BIK1 and PUB26 were used as the positive control, and PUB25 or PUB26 and GUS were used as the negative controls (Figure 2F). Taken together, these results indicate that VDAL directly interacts with the ubiquitin ligases PUB25 and PUB26 both in plant cells and *in vitro*.

### VDAL is a substrate of PUB25 and PUB26 but is not degraded *in planta*

Given that VDAL directly interacts with PUB25 and PUB26, we investigated whether VDAL is a substrate of these ubiquitin ligases. We first confirmed that the recombinant PUB25 and PUB26 proteins had Ub ligase activity (Supplemental Figures 4C and 4D). Such activity was entirely dependent on the presence of E1, E2 and one of the two E3 ligases (Supplemental Figures 4C and 4D), thus, confirming that the recombinant PUB25 and PUB26 were enzymatically active (Wang et al., 2018a; Wang et al., 2019). The addition of VDAL-HIS or GST-VDAL into the reaction led to the production of VDAL-HIS or GST-VDAL ladders, a key characteristic of ubiquitination reaction (Figures 3A and Supplemental Figure 4E). Again, the generation of ubiquitinated VDAL in this reaction required E1, E2 and PUB25 or PUB26, and reactions missing any of these components failed to ubiquitinate VDAL.

**Figure 3.**
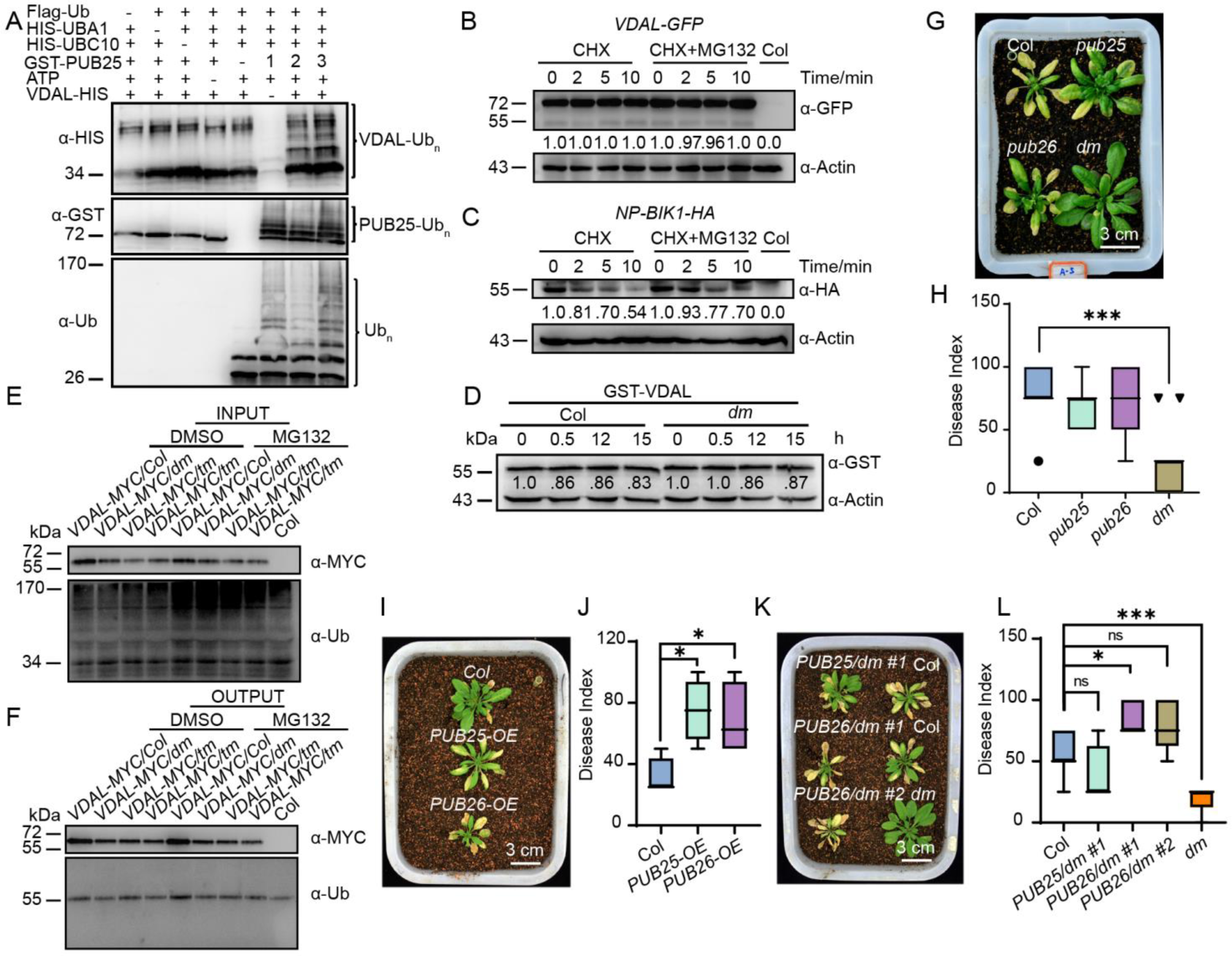
VDAL is a putative substate of PUB25 *in vitro*, and PUB25/26 negatively regulate plant resistance to Verticillium wilt. **(A)** PUB25 ubiquitinates VDAL *in vitro*. Different ubiquitination reaction systems with 40 mM ATP from different proteins purified from *E. coli* were incubated at 30 °C for 3 h. The VDAL ubiquitination and PUB25 ubiquitination were detected by anti-HIS and anti-GST antibodies, respectively. The total ubiquitination signal was detected with anti-Ub antibodies. **(B)** Addition of CHX or CHX with MG132 didn’t change the degradation pattern of VDAL. The 12-day-old *VDAL-GFP* seedlings were treated with CHX or CHX with MG132 for different times. Total proteins were extracted. Immunoblotting analysis was carried out with anti-GFP antibodies. Actin was an equal loading control. The relative protein level was quantified with ImageJ. **(C)** Addition of CHX or CHX with MG132 change the degradation pattern of BIK1. The 12-day-old *BIK1-HA* seedlings were treated with CHX or CHX with MG132 for different times. Total proteins were extracted. Immunoblotting analysis was carried out with anti-HA antibodies. Actin was an equal loading control. The relative protein level was quantified with ImageJ. **(D)** VDAL degradation has no difference between Col and *pub25 pub26* double mutant. Total proteins extracted from 12-day-old *pub25 pub26* and Col seedlings were incubated with equal GST-VDAL at 25 °C for different time. Immunoblotting analysis was carried out with anti-GST antibodies. Actin was an equal loading control. The relative protein level was quantified with ImageJ. **(E)** and **(F)** VDAL is not ubiquitinated in planta. The 12-day-old *VDAL-MYC* seedlings were treated with DMSO or MG132 for 6 hours. Total extracted proteins were immunoprecipitated with beads coated with anti-MYC antibodies. Immunoblotting analysis was carried out with anti-MYC and anti-Ub antibodies. **(G)** *pub25 pub26* double mutant exhibits more resistant to Verticillium wilt than the wild type or *pub25* and *pub26* single mutant. The plants grown in the greenhouse for two weeks were dipped into the *V. dahliae* spore suspension for 5 minutes. Photos were taken at 20 dpi. **(H)** Statistical analysis of the disease index of Col, *pub25, pub26* and *pub25 pub26* double mutant in **(G)**, count with at least 15 plants. **(I)** The PUB25 and PUB26 overexpression transgenic lines show more susceptible to Verticillium wilt caused by *V. dahliae* than the wild type. The plants grown in the greenhouse for two weeks were dipped into the *V. dahliae* spore suspension for 5 minutes. Photos were taken at 15 dpi. **(J)** Statistical analysis of the disease index of *PUB25-OE*, *PUB26-OE* and Col in **(I)**, count with at least 15 plants. **(K)** The PUB25 and PUB26 transgenic complement lines show similar phenotype with Col to Verticillium wilt caused by *V. dahliae*. The plants grown in the greenhouse for two weeks were dipped into the *V. dahliae* spore suspension for 5 minutes. Photos were taken at 20 dpi. **(L)** Statistical analysis of the disease index of plants indicated in **(K)**, count with at least 15 plants. * and *** in **(H), (J)** and **(L)** represent significant difference (P<0.05), and extremely significant difference (P<0.001) respectively, Student’s t-test. The experiments were repeated independently three times with similar results.

Next, we determined whether VDAL is degraded *in planta*. First, we treated transgenic *Arabidopsis* plants stably expressing VDAL-GFP with the protein synthesis inhibitor CHX or CHX combined with the 26S proteasome inhibitor MG132. Immunoblot analysis with anti-GFP antibodies indicated that VDAL-GFP accumulation did not change after treatment with CHX or CHX+MG132 for various periods of time (Figure 3B), but the positive control BIK1-HA is degraded (Figure 3C). To further confirm the stability of VDAL, we purified VDAL-HIS from *E. coli,* and kept VDAL solution at 25 °C for 8 days. We checked the VDAL protein by western blot for each day, and found that VDAL protein was quite stable at 25 °C, and hardly degraded (Supplemental Figure 4F). We then investigated whether PUB25 and PUB26 would modulate VDAL degradation in a cell-free assay. We purified GST-VDAL from *E. coli* and added it to total proteins extracted from wild-type or *pub25 pub26* double mutant (*dm*) plants in the presence of 10 mM ATP. We did not detect apparent differences in GST-VDAL protein levels in protein extracts from wild-type vs. *dm* plants (Figure 3D). These results suggest that VDAL is a substrate of PUB25 and PUB26, but is highly stable *in planta*. We speculated that VDAL may not be ubiquitinated in planta. We overexpressed VDAL-MYC in Col and *pub25 pub26* double mutant, and detected the ubiquitination level of VDAL in Col vs. *pub25 pub26* double mutant, to our surprise, the total ubiquitination was easily detected in input (Figure 3E), but we didn’t detect any ubiquitination of VDAL in both Col and *pub25 pub26* double mutant in output (Figure 3F). These results suggest that VDAL is not ubiquitinated and degraded by PUB25 or PUB26 *in planta*.

### PUB25 and PUB26 negatively regulate plant resistance to Verticillium wilt

Given that overexpressing VDAL increases plant resistance to Verticillium wilt and that PUB25 and PUB26 interact with VDAL, we wanted to know whether PUB25 and PUB26 are involved in plant resistance to *V. dahliae* infection. Analysis of GUS expression driven by the *PUB25/PUB26* promoters indicated that they are expressed throughout the plant (Supplemental Figures 5A and 5B). We also noticed that the subcellular localization of PUB25/PUB26-GFP fusion protein in *Arabidopsis* protoplasts was similar as VDAL-GFP (Supplemental Figures 1K, 5C-5E). We crossed the *pub25* and *pub26* single mutants obtained a *pub25 pub26* double mutant (*dm)*, and then confirmed that the expression of both genes was disrupted in the double mutant (Supplemental Figures 5F-5H). Inoculation experiments showed that the *pub25 pub26* double mutant was significantly more resistant to *V. dahliae* than *pub25* or *pub26* (Figures 3G and 3H), and that wild-type plants were more susceptible to *V. dahliae* than the *pub25 pub26* double mutants and the single mutants (Figures 3G and 3H). Under infection conditions, the disease index of *pub25 pub26* plants was significantly lower than that of both wild-type plants and the *pub25* or *pub26* single mutants (Figures 3G and 3H). Notably, the *V. dahliae* resistance of *pub25 pub26* was comparable to that of transgenic plants expressing *VDAL* (Supplemental Figures 5I and 5J).

To further explore the roles of PUB25 and PUB26 in the plant response to *V. dahliae*, we generated transgenic plants overexpressing PUB25 or PUB26 fused with Flag tag (*PUB25-OE* and *PUB26-OE* plants; Supplemental Figure 5K). After inoculation, the disease index analysis showed that the *PUB25-OE* and *PUB26-OE* plants were more susceptible to infection by *V. dahliae* than the wild type (Figures 3I and 3J). We also generated transgenic complemented lines in which the *PUB25* and *PUB26* genomic sequence was fused with *FLAG* and *MYC* tag driven by the *PUB25* or *PUB26* native promoter, respectively, in the *pub25 pub26* background (Supplemental Figures 5L and 5M). Under inoculation conditions, the complemented lines largely lost the *V. dahliae* resistance of the double mutant, as indicated by Verticillium sensitive phenotype and disease index (Figures 3K and 3L), confirming that the *V. dahliae* resistance of the double mutant was caused by *pub25 pub26* mutation. Taken together, these results indicate that PUB25 and PUB26 are negative regulators of plant immunity against infection caused by *V. dahliae*.

### PUB25 and PUB26 target MYB6 for ubiquitination and degradation

To explore the mechanism underlying the improved *V. dahliae* resistance of the *pub25 pub26* mutant, we screened an Arabidopsis yeast two-hybrid library using the ARM domain of PUB26 as bait to identify its target proteins in *Arabidopsis*. We identified one candidate interacting protein: MYB6 (Supplemental Data Set 2). The biological function of MYB6 in *Arabidopsis* has not been explored in detail. We verified the interaction of these proteins in a yeast two-hybrid assay (Figure 4A). As the C-terminus (112-237) of MYB6 had autoactivation activity in yeast harboring the AD or BD vector, we did not test it further. The full-length PUB25 and its ARM domain interacted with the N-terminus of MYB6, which harbors two MYB domains (1-111). Full-length PUB26, but not its ARM domain, interacted with the N-terminus (1-111) of MYB6 (Figure 4A). Further deletion analyses indicated that both MYB repeat domains of MYB6, including MYB6 (14-61) and MYB6 (67-111), interacted with PUB26 (Figure 4B). By contrast, no interaction was detected in yeast strains co-expressing fusion proteins with only the DNA-binding motif or the activation motif, indicating that these interactions are specific.

**Figure 4.**
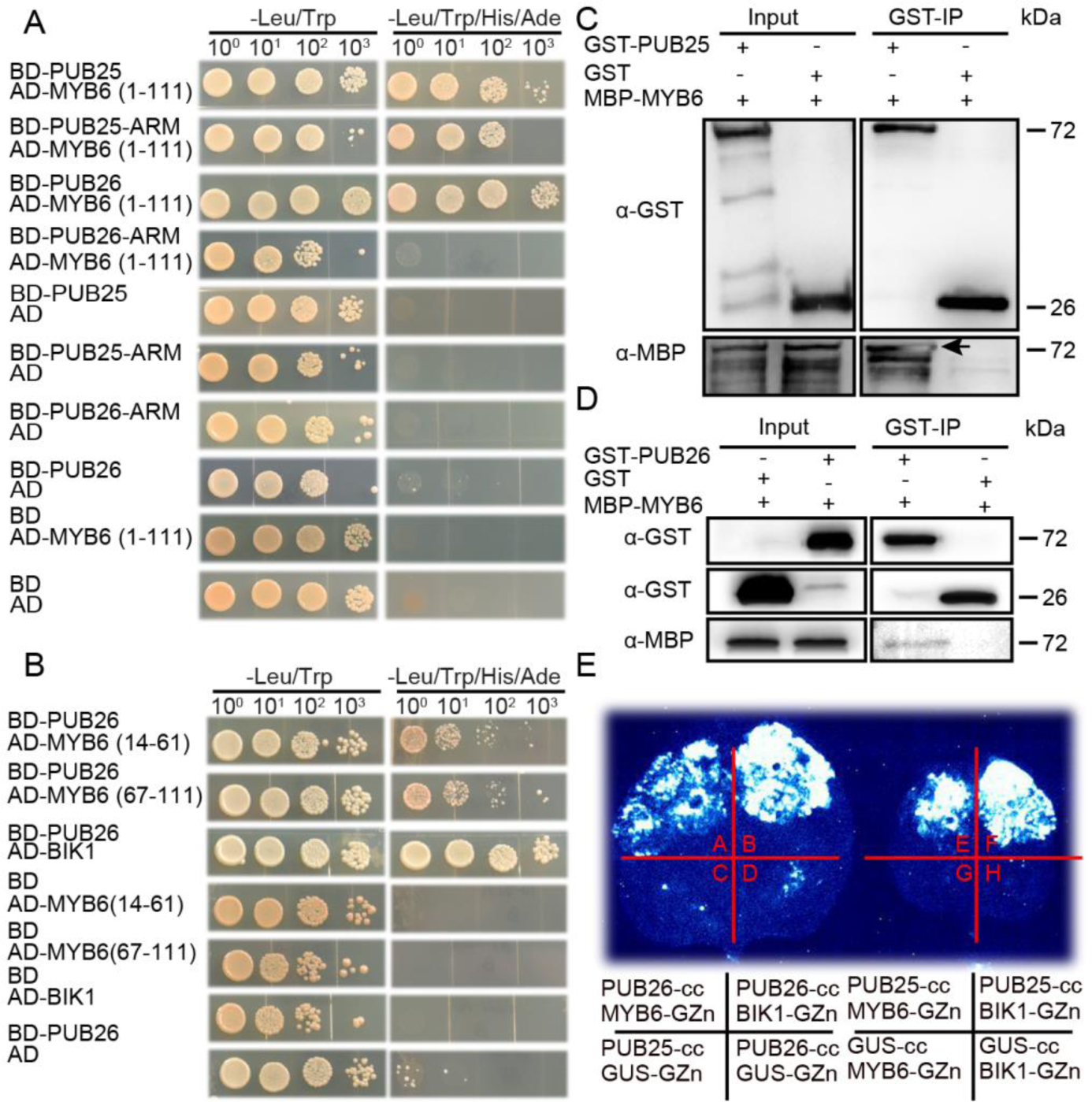
PUB25/26 interact with *Arabidopsis* transcription factor MYB6. **(A)** MYB domain (1-111) of MYB6 interacts with full length of PUB25 and PUB26, and ARM domain of PUB25 in yeast two-hybrid assay. The interaction between BD-PUB26 and AD-BIK1 is used as a positive control. Yeast cells were grown on different medium for 4 days. AD: Gal4 activation domain; BD: Gal4 DNA-binding domain. **(B)** Shorter MYB domain (14-61 and 67-111) of MYB6 interacts with full length of PUB26 in yeast two-hybrid assay. The interaction of BD-PUB26 and AD-BIK1 is a positive control. **(C)** and **(D)** MYB6 interacts with PUB25 and PUB26 in pull-down assay. Proteins were immunoprecipitated with anti-GST beads, and detected with anti-GST and anti-MBP antibodies. **(E)** MYB6 interacts with PUB25 and PUB26 in the split firefly luciferase complementation imaging (LCI) assay. Different plasmid combinations were expressed in tobacco leaves for 72 h, then images were determined by a CCD camera.

We further examined the interactions between MYB6 and the two ubiquitin ligases by performing Co-IP, pull-down and LCI assays. However, we did not detect the expression of MYB6 in *Arabidopsis* protoplasts. We then purified GST-PUB25, GST-PUB26, GST and MBP-MYB6 from *E. coli*. Pull-down assays using GST beads indicated that GST-PUB25 or GST-PUB26, but not GST, was able to pull down MBP-MYB6 (Figures 4C and 4D). Robust interactions were also observed in the LCI assay (Figure 4E), as indicated by strong luminescence only in leaf sections expressing both *MYB6* and *PUB25* or *PUB26* (Supplemental Figures 5N and 5O). These results indicate that MYB6 interacts with both ubiquitin ligases *in planta* and *in vitro*.

To determine whether MYB6 is a target of PUB25 and PUB26, we performed ubiquitination assays *in vitro* using MYB6 and both PUB25 and PUB26 isolated from *E. coli*, and found that MYB6 was ubiquitinated by both PUB25 and PUB26 (Figures 5A and Supplemental Figure 6A). Next, we investigated whether PUB25 and PUB26 modulate the degradation of MYB6 *in planta*. Analysis of transgenic Arabidopsis plants harboring the *MYB6* promoter fused to *GUS* indicated that MYB6 is widely expressed in plants (Supplemental Figure 6B). We generated transgenic *Arabidopsis* plants that stably expressed MYB6 fused with MYC, GFP or Flag tag. *MYB6* gene expression was higher in transgenic lines than in Col (Supplemental Figure 6C), but we did not detect any accumulation of MYB6 using anti-MYC, anti-GFP or anti-FLAG antibodies, respectively (Supplemental Figure 6D, only MYB-Flag is shown). However, when MYB6-Flag was expressed in the *pub25 pub26* double mutant, MYB6 was easily detected using anti-Flag antibodies (Supplemental Figure 6D). These results suggest that MYB6 is not stable in the wild type but is stable in *pub25 pub26*.

**Figure 5.**
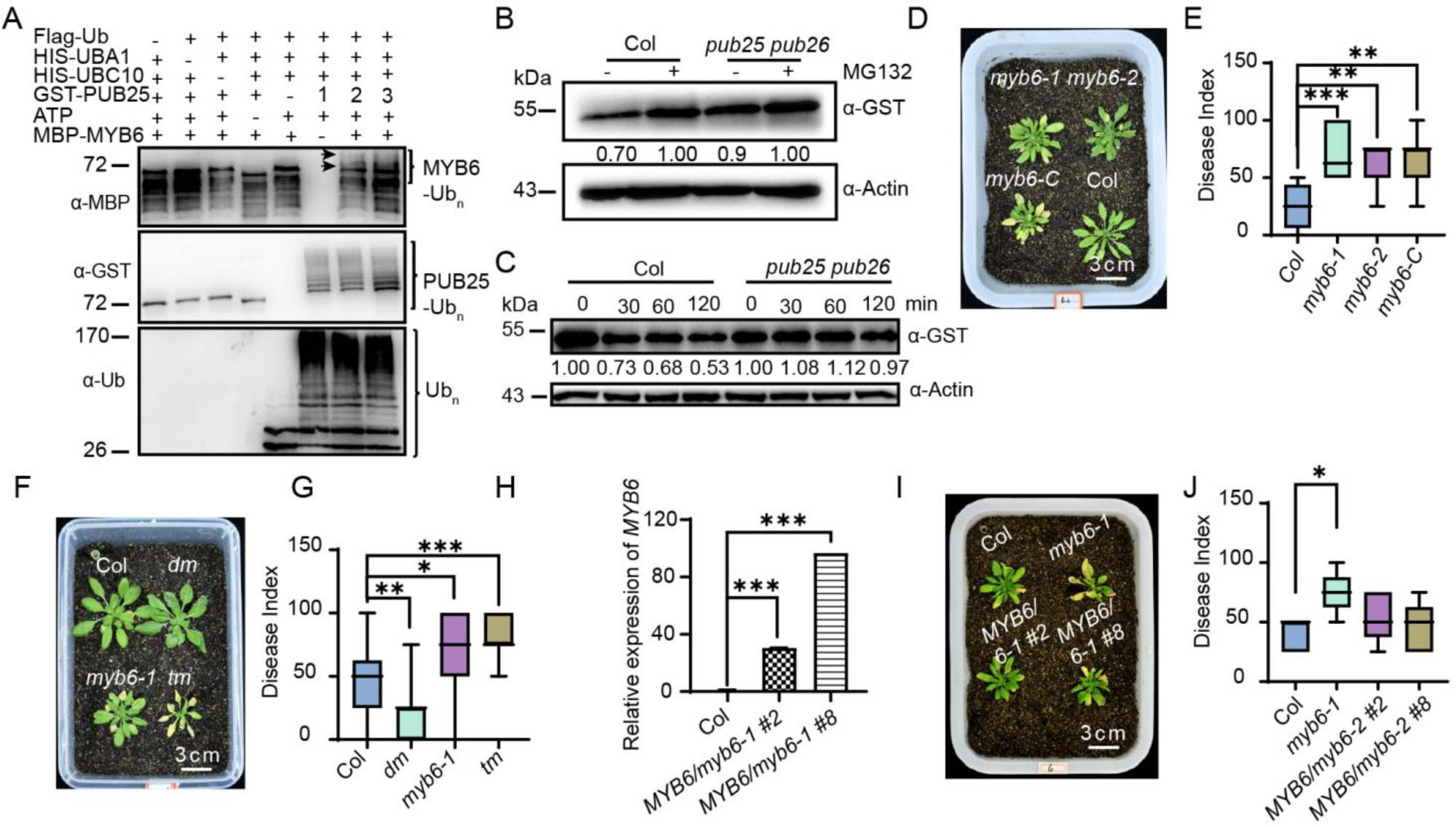
MYB6 is a target of PUB25/PUB26 and positively regulates plant resistance to Verticillium wilt. **(A)** PUB25 ubiquitinates MYB6 *in vitro*. Different ubiquitination reaction systems with 40 mM ATP composed of different proteins purified from *E. coli* were incubated at 30 °C for 3 h. The MYB6 ubiquitination and PUB25 ubiquitination were detected by anti-MBP and anti-GST antibodies, respectively. The total ubiquitination signal was detected with anti-Ub antibodies. **(B)** The 26S proteasome inhibitor MG132 treatment increases MYB6 level. Total proteins extracted from 12-day-old seedlings of Col and *pub25 pub26* were incubated with equal GST-MYB6 with or without 50 μM MG132 at 25 °C for different time. Immunoblotting analysis was carried out with anti-GST antibodies. Actin was an equal loading control. The relative protein level was quantified with ImageJ. **(C)** PUB25/26 promote MYB6 degradation. Total proteins extracted from 12-day-old Col and *pub25 pub26* seedlings were incubated with equal GST-MYB6 at 25 °C for different time. Immunoblotting analysis was carried out with anti-GST antibodies. Actin was an equal loading control. The relative protein level was quantified with ImageJ. **(D)** *myb6* mutants are more susceptible to Verticillium wilt than the wild type. The plants grown in the greenhouse for two weeks were dipped into the *V. dahliae* spore suspension for 5 minutes. Photos were taken at 15 dpi. **(E)** Statistical analysis of the disease index of Col, *myb6-1, myb6-2* and *myb6-C* in **(D)**, count with at least 15 plants. **(F)** *myb6* mutants are more susceptible to *V. dahliae*, which suppresses the *V. dahliae* resistance of *pub25 pub26* mutants. The plants grown in the greenhouse for two weeks were dipped into the *V. dahliae* spore suspension for 5 minutes. Photos were taken at 15 dpi. **(G)** Statistical analysis of the disease index of plants indicated in **(F)**, count with at least 15 plants. **(H)** The expression level of *MYB6* in their corresponding transgenic complement lines. **(I)** The MYB6 transgenic complement lines show similar disease index with Col to Verticillium wilt. The plants grown in the greenhouse for two weeks were dipped into the *V. dahliae* spore suspension for 5 minutes. Photos were taken at 15 dpi. **(J)** Statistical analysis of the disease index of plants indicated in **(I)**, count with at least 15 plants. *, ** and *** in **(E), (G), (H)** and **(J)** represent significant difference (P<0.05), highly significant difference (P<0.01) and extremely significant difference (P<0.001) respectively, Student’s t-test. The experiments were repeated independently three times with similar results.

We then performed a cell-free assay to determine whether the stability of MYB6 is regulated by PUB25 and PUB26. First, we added GST-MYB6 into cell extracts from wild type and *pub25 pub26* plants with or without the 26S proteasome inhibitor MG132 and observed that MG132 inhibited the degradation of GST-MYB6 in the presence of ATP (Figure 5B). We then incubated purified GST-MYB6 with total proteins extracted from wild-type or *pub25 pub26* plants in the presence of 10 mM ATP for different periods of time and detected changes in protein levels. GST-MYB6 degraded much more slowly in *pub25 pub26* vs. the wild type (Figure 5C). These results suggest that PUB25 and PUB26 modulate the stability of MYB6 via the 26S proteasome.

### *myb6* is susceptible to and suppresses the resistance of *pub25 pub26 to V. dahliae*

The finding that MYB6 is ubiquitinated by PUB25 and PUB26 suggests that it is involved in plant resistance to *V. dahliae*. We examined this hypothesis using an inoculation experiment. We obtained three *MYB6* mutants (*myb6-1, myb6-2* and *myb6-Cas9*) (Supplemental Figures 6E-6H) and the *pub25 pub26 myb6-1* triple mutant (*tm*) (Supplemental Figure 6I). Under normal growth conditions, the mutants did not exhibit any discernible phenotypes in terms of growth, development or reproduction compared to the wild type. However, under *V. dahliae* infection, the three *myb6* mutants were more susceptible to *V. dahliae* infection than the wild type (Figures 5D and 5E) and the *pub25 pub26* double mutant (Figures 5F and 5G). The *V. dahliae* susceptibility of the *myb6 pub25 pub26* triple mutant was similar to that of *myb6* (Figures 5 and 5G). Under our experimental conditions, the disease index of the *myb6 pub25 pub26* triple mutant was approximately 75, which is similar to the disease index (65 to 75) of the *myb6* mutants, whereas the disease indices of the wild type and *pub25 pub26* were approximately 25 and 50, respectively (Figures 5F and 5G). *pub25 pub26* overexpressing *MYB6-Flag* showed similar or slightly higher levels of disease resistance than *pub25 pub26*, which was more resistant than the wild type (Supplemental Figures 6J and 6K). These results suggest that *MYB6* acts at downstream of *PUB25* and *PUB26* in response to *V. dahliae* infection.

**Figure 6.**
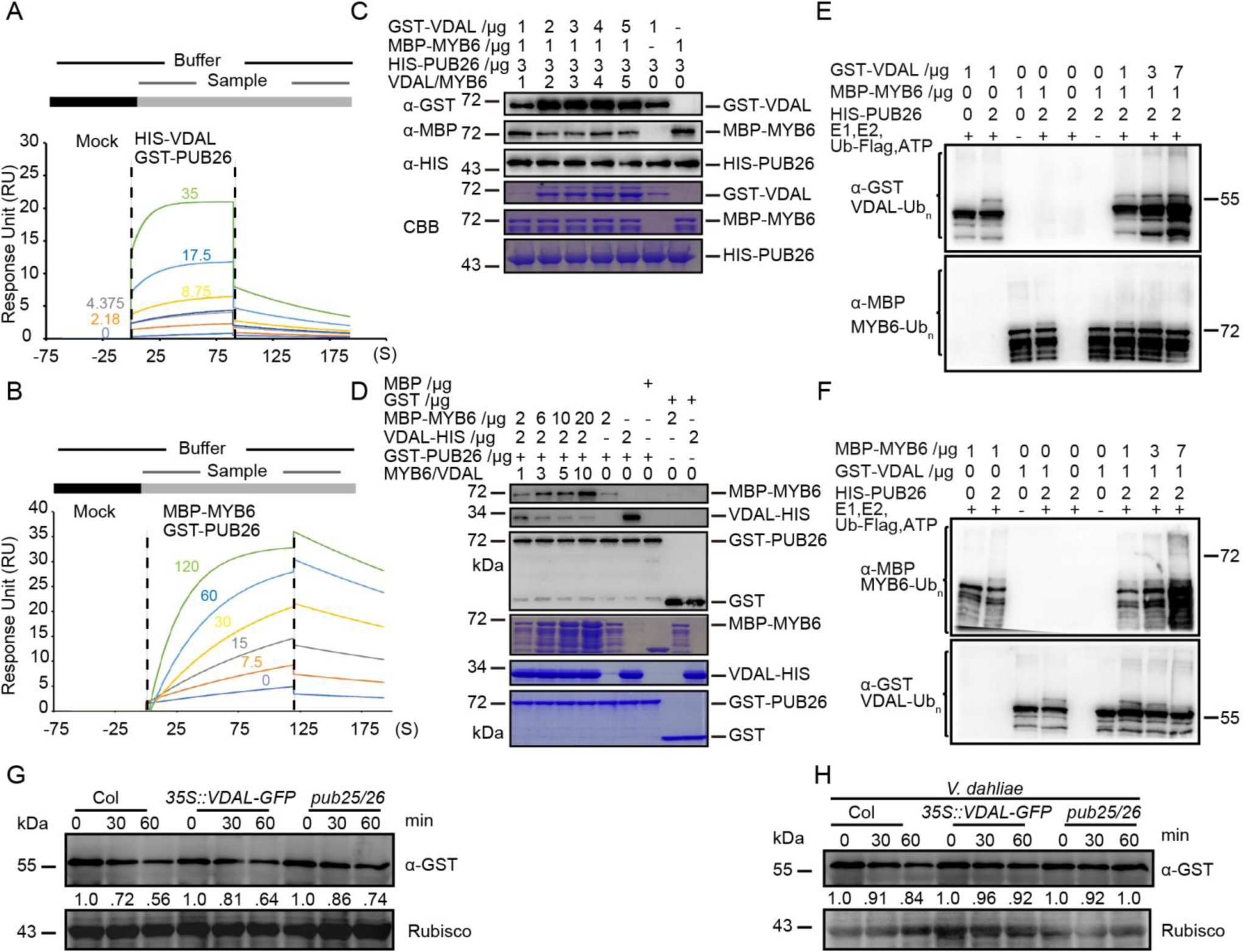
VDAL and MYB6 are competitive in interacting with PUB26. **(A)** and **(B)** Affinity characterization of HIS-VDAL **(A)** or MBP-MYB6 **(B)** to GST-PUB26. The anti-GST antibodies were immobilized onto CM5 chip. After baseline was established, the first sample GST-PUB26 injected over the chip was followed by the second sample HIS-VDAL **(A)** or MBP-MYB6 **(B)** for certain time with different concentration. The signals generated were subtracted from reference channel and then analyzed with kinetics fitting. The numbers represent the different concentration (ug/ml) of the second sample. **(C)** VDAL inhibits the interaction between PUB26 and MYB6 in the competitive pull down assay. First, HIS-PUB26 was immunoprecipitated with anti-HIS beads, then GST-VDAL and MBP-MYB6 were added into reaction together with different concentration for 2 h. After fully washing for non-interaction proteins, proteins from beads were detected with anti-GST, anti-HIS and anti-MBP antibodies. **(D)** MYB6 inhibits the interaction between PUB26 and VDAL in a competitive pull-down assay. First, GST-PUB26 was immunoprecipitated with anti-GST beads, then VDAL-HIS and MBP-MYB6 were added into reaction with different concentration. Pull-down proteins were detected with anti-GST, anti-HIS and anti-MBP antibodies. **(E)** and **(F)** VDAL and MYB6 are competitive to be ubiquitinated by PUB26. Different ubiquitination reaction systems with 40 mM ATP composed of different proteins purified from *E. coli* were incubated at 30 °C for 3 h. The ubiquitinated VDAL and MYB6 were detected by anti-GST or anti-MBP anibodies. The total ubiquitination signal was detected with anti-Ub antibodies. **(G)** and **(H)** Comparison of MYB6 degradation rate among Col, *VDAL-OE* and *pub25 pub26* double mutant. Total proteins extracted from12-day-old Col, *VDAL-OE* and *pub25 pub26* seedlings treated with or without *V. dahliae* for 12 h were incubated with equal GST-MYB6 at 25 °C for different time. Immunoblotting analysis was carried out with anti-GST antibodies. Actin was an equal loading control. The relative protein level was quantified with ImageJ.

To confirm that the enhanced susceptibility of the *myb6* mutants to *V. dahliae* infection was caused by *MYB6* mutation, we introduced *MYB6* genomic DNA driven by the *MYB6* promoter into the *myb6-1* mutant background (Figure 5H). Inoculation experiments showed that two transgenic *myb6* lines had comparable or even a little higher level of disease resistance than the wild type (Figures 5I and 5J). These results suggest that *MYB6* positively regulates plant resistance to wilt caused by *V. dahliae* and acts at downstream of *PUB25* and *PUB26*.

### VDAL competes with MYB6 for interactions with PUB25 and PUB26

Given that both VDAL and MYB6 interact with PUB25 and PUB26, we explored why overexpressing VDAL increased plant resistance to *V. dahliae* infection. We hypothesized that VDAL might compete with MYB6 for interacting with PUB25 and PUB26, thus hijacking these proteins to reduce the ubiquitination and degradation of MYB6. We performed the Biacore assay to measure the affinity of HIS-VDAL and MBP-MYB6 for GST-PUB26 using proteins purified from *E. coli*. The anti-GST antibodies were immobilized onto a CM5 chip. After the baseline was established, the first sample (GST-PUB26 in Figures 6A, 6B and Supplemental Figure 7B, GST in Supplemental Figure 7A) was overlaid on the chip, followed by the second sample (HIS-VDAL in Figures 6A and Supplemental Figure 7A, MBP-MYB6 in Figures 6B and MBP in Supplemental Figure 7B), and incubated for various periods of time. Strong interaction signals were detected between HIS-VDAL and GST-PUB26, between MBP-MYB6 and GST-PUB26. The interaction signal Response Unit (RU) values between HIS-VDAL and GST-PUB26 and between MBP-MYB6 and GST-PUB26 were comparable (Figure 6A and 6B). However, the association rate (ka in Supplemental Table 1), disassociation rate (kd in Supplemental Table 1) and affinity (KD in Supplemental Table 1) of HIS-VDAL and GST-PUB26 were higher, suggesting that HIS-VDAL and GST-PUB26 associate more rapidly and have a stronger affinity than MBP-MYB6 and GST-PUB26. As expected, no interaction signals were detected between HIS-VDAL and GST or between GST-PUB26 and MBP (Supplemental Figures 7A and 7B, Supplemental Table 1). These results suggest that the association between VDAL and PUB25 or PUB26 may leads to the occupation of the PUBs, thereby providing favorable conditions for the accumulation of MYB6 to help the plant resist infection by *V. dahliae.* Similar results were obtained in a subsequent competitive pull-down assay. The interaction between MBP-MYB6 and HIS-PUB26 decreased with increasing amounts of GST-VDAL (Figures 6C) and the interaction between VDAL-HIS and GST-PUB26 decreased with increasing amounts of MBP-MYB6 (Figures 6D).

**Figure 7.**
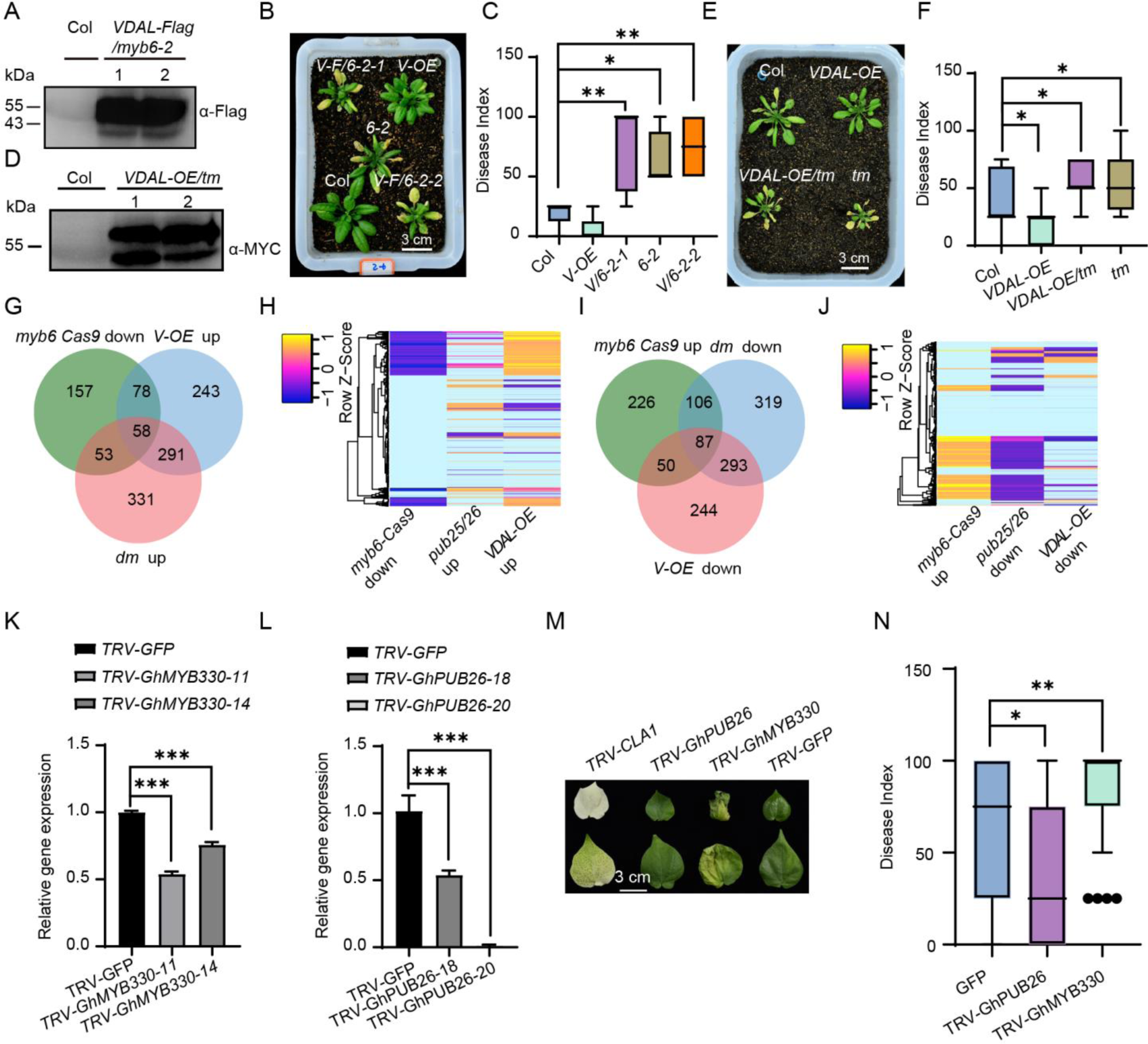
VDAL competes with MYB6 for PUB25/26-mediated degradation in plants. **(A)** The accumulation of VDAL protein in two *myb6-2* transgenic lines. Total proteins extracted from 12-day-old Col and *VDAL-Flag/myb6-2* seedlings were detected with anti-Flag antibodies. **(B)** The phenotype comparison of disease symptoms in the wild type (Col), *VDAL* overexpression line (Col, *V-OE*), *myb6-2* (6-2), *myb6-2* overexpressing VDAL line 1 and 2 (*V/6-2-1*, *V/6-2-2*). Photos were taken at 15 dpi. **(C)** Statistical analysis of the disease index of plants indicated in **(B)**, count with at least 15 plants. **(D)** VDAL-MYC overexpression in transgenic *pub25 pub26 myb6-1* (*tm*) triple mutants. Total proteins extracted from 12-day-old Col and *VDAL-MYC /pub25 pub26 myb6-1* seedlings were detected with anti-MYC antibodies. **(E)** The susceptibility of *VDAL-OE/tm* to *V. dahliae* is similar as *myb6*, and more than Col. The plants grown in the greenhouse for two weeks were dipped into the *V. dahliae* spore suspension for 5 minutes. Photos were taken at 20 dpi. **(F)** Statistical analysis of the disease index of plants indicated in **(E)**, count with at least 15 plants. **(G-J)** Transcriptome analysis of the *myb6-Cas9, VDAL-OE* and *pub25/26* double mutant by RNA-Seq after treatment with *V. dahliae* spores (1×10^6^ conidia/mL) for 12 h. Data were obtained from three independent experiments. **(G)** and **(I)** Venn diagrams showing the differentially regulated genes (down- and upregulated genes) overlapped between the different genotype. (H) and (J) The heat map was drawn according to the expression levels in the *myb6-Cas9*, *VDAL-OE* and *pub25/26* compared with Col. **(K)** The relative expression level of *GhMYB330* in the VIGS cotton. **(L)** The relative expression level of *GhPUB26* in the VIGS cotton. **(M)** and **(N)** The knock down of *GhMYB330* causes cotton susceptible to *V. dahliae* and knock down of *GhPUB26* causes cotton resistance to *V. dahliae.* The plants grown in the greenhouse for three weeks were injected 2 mL *V. dahliae* spore suspension (1×10^6^ conidia/mL) through stem near cotyledon. Photos were taken at 20 dpi, count with at least 25 plants. * and ** in **(C), (F), (K), (L)** and **(N)** represents significant difference (P<0.05) and highly significant difference (P<0.01), Student’s t-test. The experiments were repeated independently three times with similar results.

Given that VDAL competes with MYB6 for binding to PUBs, we performed a competitive ubiquitination in vitro assay using purified VDAL-HIS, GST-PUB25, HIS-PUB26 and MBP-MYB6 proteins. The ubiquitination level of MBP-MYB6 decreased with increasing amounts of VDAL (Figures 6E and Supplemental Figure 7C) and vice versa (Figures 6F and Supplemental Figure 7D).

Next, we performed a cell-free assay using *Arabidopsis* seedlings with or without *V. dahliae* infection to measure the degradation status of MYB6. We added GST-MYB6 protein into cell extracts from wild-type, *VDAL-OE* and *pub25 pub26* seedlings with or without *V. dahliae* infection in the presence of ATP for different periods of time. GST-MYB6 degraded at a similar rate or slightly more rapidly in the *VDAL-OE* line compared to *pub25 pub26*, but at the fastest speed in the wild type in both the presence and absence of *V. dahliae* infection (Figures 6G and 6H). These results suggest that VDAL competes with MYB6 for interactions with PUB25 and PUB26. Therefore, MYB6 might escape from PUB25- and PUB26-mediated degradation to enhance plant resistance to *V. dahliae* infection *in planta*.

To further confirm that the function of VDAL depends on MYB6 in *Arabidopsis*, we overexpressed *VDAL* fused with a Flag tag in *myb6-2* (Figures 7A) and compared the disease resistance of these plants vs. *myb6-2*. Overexpressing *VDAL* did not increase the resistance of *myb6-2* to *V. dahliae* infection (Figures 7B-7C). To further verify this notion *in planta*, we generated transgenic lines stably expressing *VDAL-MYC* in the *myb6 pub25 pub26* triple mutant background (*VDAL-OE/tm*) to examine its potential role in triggering plant immune responses (Figures 7D). After inoculation, *VDAL-OE/tm* exhibited similar levels of susceptibility to *V. dahliae* to the *pub25 pub26 myb6-1* triple mutant, as indicated by disease index (Figures 7F).

These results suggest that VDAL itself does not have a direct effect on increasing resistance to *V. dahliae* infection; this effect likely depends on MYB6.

### VDAL and PUB25/PUB26 regulate plant resistance to wilt caused by *V. dahliae*

As both *pub25 pub26* plants and plants stably expressing VDAL exhibited similar levels of resistance to *V. dahliae* infection, we reasoned that VDAL and PUB25/PUB26 might function via a similar molecular mechanism. More importantly, we demonstrated that MYB6 can escape from PUB25- and PUB26-mediated degradation to enhance plant resistance to *V. dahliae* infection in *VDAL* overexpression plants. These findings prompted us to explore the transcriptome profiles of plants with various genotypes. We treated ten-day-old seedlings with or without 10^6^ *V. dahliae* spores for 12 h, isolated total RNAs from the samples, and subjected them to RNA-deep sequencing on the Illumina HiSeq platform. We generated 150-bp high-quality trimmed paired-end reads and mapped them to the *Arabidopsis* genome (TAIR10) using bowtie2 with default settings. We collected raw counts from three independent experiments (each sample with 6.0 G clean data), normalized them and performed differential gene expression analyses in Col, *myb6-Cas9*, *pub25 pub26* or *VDAL-OE* lines under control condition (water treatment) and 12 h *V. dahliae* treatment using edgeR, respectively. The genes significantly induced by *V. dahliae* in the Col group were chosen for the comparison with the expression levels of the treatment samples between different groups. The relative expression level of a gene was defined as the fold change of the gene in *V. dahliae*-treated *myb6-Cas9*, *pub25 pub26* or *VDAL-OE* samples minus the fold change of the same gene in *V. dahliae*-treated Col samples, respectively. Thus positive or negative value of the relative expression level indicated the change level of the gene in *myb6-Cas9*, *pub25 pub26* or *VDAL-OE* was higher or lower than the change level in Col. We constructed Venn diagrams showing the expression patterns of common differentially expressed genes (DEGs) in different samples using the online tool jvenn (http://jvenn.toulouse.inra.fr/app/example.html).

We identified 2196 differentially expressed genes (DEG) in Col of *V. dahliae* treatment compared to water treatment group (using *P*<0.05, t-test and |logFC| >1) (Supplemental Data Set 3). Among these DEGs, 469, 670 and 733 genes had higher relative expression levels, while 346, 674 and 805 genes had lower expression levels in *myb6-Cas9*, *VDAL-OE* and *pub25 pub26* plants than Col, respectively (Figures 7G and 7H). Of the overlapping DEGs, 349 genes were higher in both *VDAL-OE* and *pub25 pub26*, comprising 52% of up-regulated DEGs in *VDAL-OE* and 48% in *pub25 pub26*; 136 genes were up-regulated in *VDAL-OE* and down-regulated in *myb6-Cas9*, comprising 64.7% of down-regulated DEGs in *myb6-Cas9*; 111 genes were up-regulated in *pub25 pub26* and down-regulated in *myb6-Cas9*, comprising 47.2% of down-regulated DEGs in *myb6-Cas9*; 58 genes were down-regulated in *myb6-Cas9* and up-regulated in both *VDAL-OE* and *pub25 pub26* (Figure 7G, and 7H for heatmap, Supplemental Data Set 4). Similarly, 380 genes were down-regulated in both *VDAL-OE* and *pub25 pub26*, and 193 DEGs were up-regulated in *myb6-Cas9* and down-regulated in *pub25 pub26*; 137 gene were up-regulated in *myb6-Cas9* and down-regulated in *VDAL-OE*; and 87 genes were up-regulated in *myb6-Cas9* and down-regulated in *VDAL-OE* and *pub25 pub26* (Figure 7I, and 7J for heatmap, Supplemental Data Set 5). These DEGs belong to different categories. 58 overlapped genes in *myb6* down-regulated and *pub25 pub26* and *VDAL-OE* up-regulated genes are highly enriched in response to stimulus and immune process (Supplemental Figure 8A), and 87 overlapped genes in *myb6* up-regulated and *pub25/26* and *VDAL-OE* down-regulated genes are highly enriched in response to stimulus (Supplemental Figure 8B). However, we did not find any difference in chitin-activated MPK3, 4 and 6 (Wan et al., 2004) (Supplemental Figures 8C-8E), and the expression of the PAMP-responsive gene *ARABIDOPSIS NON-RACE SPECIFIC DISEASE RESISTANCE GENE (NDR1)/HAIRPIN-INDUCED GENE (HIN1)-LIKE 10* (*NHL10*, a marker gene in the MPK-mediated defense pathway) (Sheikh et al., 2016) (Supplemental Figure 8F), the PTI marker gene *FLG22-INDUCED RECEPTOR-LIKE KINASE 1* (*FRK1*, *At2g19190*) (Asai et al., 2002) (Supplemental Figure 8G), the SA marker gene *PATHOGENESIS RELATED 1* (*PR1*) (Uknes et al., 1992) did not show apparent difference under *V. dahliae* treatment among different plants (Supplemental Figure 8H).

**Figure 8.**
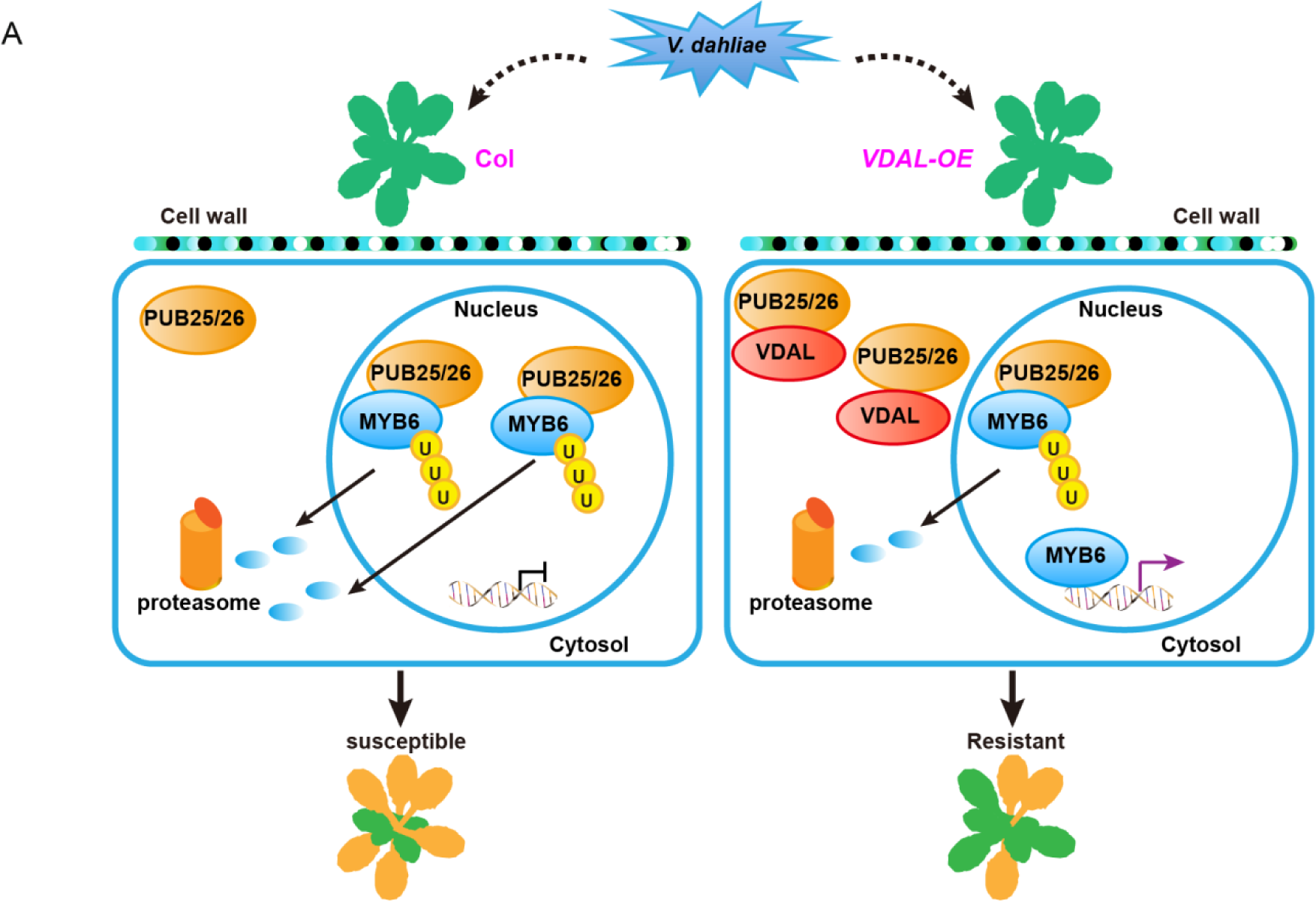
(A) A proposed model for VDAL in increasing resistance to *V. dahliae* via interacting with E3 ligases PUB25 and PUB26. In wild-type plants, PUB25 and PUB26 ubiquitinate the transcription factor MYB6, and mediate its degradation through the 26S proteasome. In VDAL overexpressing plants, VDAL competitively binds PUB25 and PUB26, which results in reducing the degradation of MYB6. As a positive regulator, the accumulated MYB6 increases disease resistance.

These results indicate that VDAL, PUB25 and PUB26 are involved in plant resistance to *V. dahliae* by functioning in the similar immunity pathways that are different from chitin-, Flg22- or SA-mediated pathways (Wang et al., 2020b), but they also individually regulate some other genes. Our results also suggest that the enhanced resistance of *pub25 pub26* to *V. dahliae* is likely due to the accumulation of MYB6.

Given that both the transgenic cotton and *Arabidopsis* enhanced resistance to *V. dahliae* infection, it promoted us to explore whether this mechanism was similar or not between these two plants. We silenced the homologous genes (*GhPUB26* and *GhMYB330*) of *Arabidopsis PUB26* and *MYB6* in island cotton by Virus-Induced Gene Silencing (VIGS) (Sarde et al., 2019) (Figure 7K and 7L). Here *chloroplastos alterados 1* (*CLA1*) was used as a positive reporter control for silencing efficiency. The inoculation experiment showed that the knock down of *GhPUB26* caused cotton more resistance to *V. dahliae*, while the knock down of *GhMYB330* caused cotton more susceptible to *V. dahliae* (Figure 7M and 7N) compared with GFP control. These results suggest that the transgenic cotton plants overexpressing *VDAL* have a similar disease resistance mechanism to *V. dahliae* as the *VDAL* transgenic *Arabidopsis* plants.

## DISCUSSION

The soil-borne hemibiotrophic fungal pathogen *V. dahliae* secretes more than 700 proteins (Klosterman et al., 2011), but only a few have been identified as effectors, such as VdSCP41 (Qin et al., 2018), AVe1 (Fradin et al., 2009; Deng et al., 2015), VdIsc1 (Liu et al., 2014a) and VdSCP7 (Zhang et al., 2017) mentioned in the introduction. In this study, we demonstrated that VDAL is a secretory elicitor protein that can cause leaf wilting and ROS burst when the protein was inoculated in leaves, which is very similar to bacterial Harpins (Choi et al., 2013). Some effectors produced by hemibiotrophic fungi such as *V. dahliae* (such as VdSCP41 and VdIsc1) inhibit plant immunity, whereas others (such as AVe1 and VDAL) increase plant immunity when over-expressed in plant cells (Castroverde et al., 2016). It was found that overexpressing *Ave1* in Ve1(-) tomato induced the expression of defense genes, suggesting that Ave1 in the cytosol triggers plant immunity independently of the Ve1 receptor (Castroverde et al., 2016). The current findings suggest that VDAL can increase the Verticillium wilt resistance when overexpressed in both cotton and *Arabidopsis*.

Previous studies have found that several effectors from different pathogens can directly hijack host 26S proteasomes, thus inhibiting the degradation of immune-related proteins for their successful infection. For example, HopM1 from *Pseudomonas syringae* destabilizes AtMINs (*Arabidopsis thaliana* HopM interactors, belonging to the ARF GEF proteins) in a proteasome-dependent manner (Nomura et al., 2006; Nomura et al., 2011). However, HopM1, HopAO1, HopA1, and HopG1 were found to be the putative proteasome inhibitors, which makes it a little more complex to explain the function of HopM1 for promoting degradation of AtMINs (Ustun et al., 2016). HopM1 can directly interact with some E3 ubiquitin ligases and proteasome subunits, suggesting its roles as a potential proteasome inhibitor (Ustun et al., 2016). *Phytophthora infestans* effector AVR3a stabilizes potato U-box E3 ligase CMPG1, and suppresses the degradation of CMPG1, thus preventing infestin 1-mediated host cell death during the biotrophic phase (Ustun et al., 2016). The rice blast fungus *Magnaporthe oryzae* effector AvrPiz-t inhibits the rice RING E3 ubiquitin ligase APIP6 (AvrPiz-t Interacting Protein 6) for enhancing the susceptibility of rice to *M. oryzae* (Park et al., 2012). Interestingly, APIP6 ubiquitinates AvrPiz-t *in vitro*, and both APIP6 and AvrPiz-t are degraded when coexpressed in the transient assays (Park et al., 2012). Furthermore, the effector AvrPtoB is an E3 ligase that can use the host 26S proteasome to directly target several host immunity-related proteins such as the SA receptor NPR1 (Janjusevic et al., 2006; Chen et al., 2017), tomato protein kinase Fen (Rosebrock et al., 2007; Ntoukakis et al., 2009), FLS2 (Goehre et al., 2008), CERK1 (Gimenez-Ibanez et al., 2009) for their degradation to subvert plant immunity.

Different from the abovementioned effectors that mainly target the host E3 ligases or they are E3 ligases targeting host proteins for disturbing the plant immunity, our results indicate that VDAL targets and interferes with two negative E3 ligases PUB25 and PUB26 in plant immunity, but is not degraded *in vivo,* and also very stable *in vitro*. Cell free assay indicated that there is no significant difference in the degradation rate of VDAL protein in Col and *pub25 pub26* double mutant. We found that the localization pattern of VDAL-GFP in protoplasts and transgenic plants were similar with that of PUB25/26-GFP in protoplasts, which provided a basis for them to work together. Plants overexpressing *VDAL* and the *pub25 pub26* double mutant showed similar resistance to *V. dahliae* infection, i.e., more resistant than the wild type, suggesting that they genetically function in the same pathway. Transgenic plants overexpressing either *PUB25* or *PUB26* were more susceptible to *V. dahliae* infection than wild-type plants. These findings indicate that PUB25 and PUB26 play negative roles in plant resistance to *V. dahliae*, which are similar as their roles in response to the fungal pathogen *Botrytis cinerea* and the non-virulent bacterial strain *Pseudomonas syringae* pv. *tomato (Pto*) DC3000 *hrcC*^−^ (Wang et al., 2018a).

Furthermore, we found that VDAL is not ubiquitinated in plants, but could be ubiquitinated *in vitro*, suggesting that VDAL mainly hijacks PUBs to prevent their roles for degradation of MYB6, which leads to MYB6 protein accumulation and disease resistance (Figure 8). One possibility is that VDAL is localized in some unknown big granule structures that may protect VDAL from degradation by PUB25/26. This phenomenon needs further explored in the future. As PUB25/26 also target the positive regulator of BIK1 in plant immune response for its degradation (Wang et al., 2018a), we also speculate that the BIK1 would be less affected by PUB25/26 in VDAL transgenic plants, thus increasing disease resistance, which needs further explored in the future study. Genetics analysis indicated that *myb6* mutants were more susceptible to *V. dahliae* than the wild type, while the *pub25 pub26 myb6* triple mutant showed a similar susceptibility to *V. dahliae* as the *myb6* single mutant. The *pub25 pub26* double mutant overexpressing *MYB6* was more resistant to *V. dahliae* than the wild type. MYB6 protein is hard to be detected in the wild type overexpressing MYB6, but accumulated to a high level in *pub25 pub26* mutant. However, the disease resistance is comparable between the *pub25 pub26* and wild type overexpressing *MYB6*, implying that the presence of MYB6 is necessary, but not sufficient for increasing disease resistance. These findings indicate that PUB25 and PUB26 directly target MYB6 and negatively regulate MYB6 stability during the pathogen response, and MYB6 is a positive regulator in resistance for *V. dahliae* disease. Notably, plants overexpressing VDAL in the *myb6* mutant showed similar susceptibility to *V. dahliae* as the *myb6* single mutant, suggesting that increasing resistance to *V. dahliae* infection by overexpressing *VDAL* depends on MYB6. Our Biacore and competitive pull-down assays suggested that VDAL interacts with PUB25 and PUB26, thus interfering with their interactions with MYB6, and reducing MYB6 degradation, thereby leading to increased resistance to *V. dahliae* infection. In the current study, our results showed that VDAL possess a signal peptide that has secreting activity. This is consistent with the notion that effectors are characterized by N-terminal signal peptide which can lead them to the secret pathway (Sperschneider et al., 2017; Wang et al., 2021). The VDAL homologs widely exist in other fungi. We speculated that VDAL is secreted into the extracellular space as an elicitor to attack plants, or even into host cells. There are two possible explanations for VDAL roles inside host cells: one explanation is that as a hemibiotrophic fungal pathogen, *V. dahliae* might use VDAL to target PUB25/26 in order to accumulate more MYB6, thus not killing the plant cells immediately during its infection; or plants evolve a new mechanism to exploit VDAL to fight against *V. dahliae*.

We determined the kinase activity of MAPK3/4/6 in the *VDAL* transgenic plants and *myb6* mutant after treatment with chitin or *V. dahliae*, and did not find any obvious difference. The transcripts by qRT-PCR analyses indicate that the expression of some marker genes such as the MPK-mediated marker gene *NHL10* (Sheikh et al., 2016), the PTI marker gene *FRK1* (Asai et al., 2002) and the SA marker gene *PR1* (Uknes et al., 1992) were not changed in the *VDAL* transgenic plants and *myb6* mutants. However, we found that the expression of some disease-related genes encoding such as TIR-NBS-LRR protein and WRKY transcriptional factors (please see supplemental data set 4-5) was changed in VDAL plants and *myb6* mutants. These results suggest that MYB6-mediated plant immune response is not through these well studied traditional pathways such as the systemic acquired resistance and HR during bacterial infection (Wang et al., 2020b; Zhou and Zhang, 2020). However, we cannot exclude the possible role of BIK1 in plant resistance to Verticillium wilt as BIK1 is targeted by PUB25/26 (Wang et al., 2018a), and BIK1 plays critical roles in both ETI and PTI for plant resistance to *P. syringae pv. tomato* DC3000 strain (Ngou et al., 2021; Yuan et al., 2021b; Yuan et al., 2021a).

When plants are infected by pathogens, plant growth is usually retarded due to the growth-defense trade-off. In this study, *Arabidopsis* and cotton plants overexpressing *VDAL* did not show any growth or developmental defects, but they showed increased resistance to *V. dahliae*. We found that GhMYB330 and GhPUB26 in cotton have the similar physiological function to *V. dahliae* infection as MYB6 and PUB26 in *Arabidopsis*, which suggests that the mechanism of plant disease resistance caused by overexpression of VDAL is conserved. A recent study involving the successful cloning of the glutathione S-transferase gene *Fhb7* from *Triticum aestivum* indicated that the horizontal transfer of this gene from a fungus underlines *Fusarium* head blight resistance in wheat (Wang et al., 2020a). Therefore, *VDAL* represents a good candidate gene from *V. dahliae* for improving plant resistance to *V. dahliae* infection without yield penalty when overexpressed in plants. Although many effector proteins such as Harpins (Peng et al., 2004), SSB_Xoc_ (Cao et al., 2018), PsCRN161 (Rajput et al., 2015) are found to increase the disease resistance when expressed in plants without affecting plant development and growth, the molecular mechanisms are not well understood. Thus, our results provide a molecular basis to highlight the possibility for using genes from the pathogens to improve crop pathogen resistance.

## METHODS

### Plant Materials and Growth Conditions

All transgenic *Arabidopsis thaliana* lines and T-DNA insertion mutants used in this study are in the Columbia-0 (Col-0) ecotype background. The T-DNA insertion mutants *pub25* (AT3G19380, SALK-147032C), *pub26* (AT1G49780, CS351943,GK308D07), *myb6-1* (AT4G09460, SALK_074789C) and *myb6-2* (AT4G09460, CS403018) were obtained from the ABRC (*Arabidopsis* Biological Resource Center). Homozygous lines were obtained by PCR using T-DNA specific and the gene-specific primers. *Pro35S::VDAL-GFP*, *Pro35S::VDAL-MYC, Pro35S::MYB6-Flag*, *Pro35S::PUB25-GFP*, and *Pro35S::PUB26-GFP* were introduced into Col by the floral-dip method (Clough and Bent, 1998) to generate overexpression transgenic lines. For complement lines of *myb6-1*, *MYB6* genomic DNA including the native promoter and coding region were amplified and cloned into PCAMBIA1301 vector (*NP::MYB6-eGFP*), then transferred into *myb6-1* mutant. *Pro35S::PUB25-Flag*, *Pro35S::PUB26-Flag*, *ProPUB25::PUB25-Flag* (Wang et al., 2018a) and *ProPUB26::PUB26-MYC* were described previously (Wang et al., 2019) *Arabidopsis* seeds were sown on Murashige and Skoog (MS) medium containing 2% sucrose and 0.8% agar. 7- or 10-days seedlings were transferred to soil and grown under short-day (12-h light/12-h dark) or long-day (16-h light/8-h dark) conditions in a growth room at 20-22°C.

### Vector Construction

For constructing *Pro35S::VDAL-GFP* and *Pro35S::VDAL-MYC*, the cDNA of *V. dahliae VDAL* was amplified and cloned into pCAMBIA1300 vector driven by *35S* promoter. For *Pro35S::MYB6-MYC/Flag/GFP*, *Pro35S::PUB25-GFP* and *Pro35S::PUB26-GFP*, the cDNA of *MYB6, PUB25* or *PUB26* from Col was cloned into pCAMBIA1300 vector driven by *35S* promoter and fused to *MYC*, *Flag* or *GFP* tag, respectively. For *ProMYB6::MYB6-eGFP*, the promoter and genomic DNA of *MYB6* from Col genomic DNA were cloned into pCAMBIA1301 vector and fused to *GFP* tag. For *Pro35S::PUB25-ccLUC*, *Pro35S::PUB26-ccLUC*, *Pro35S::MYB6-GZnLUC*, *Pro35S::VDAL-GZnLUC*, *Pro35S::BIK1-GZnLUC*, *Pro35S::GUS-GZnLUC*, *Pro35S:: GUS-ccLUC*, the CDS of *PUB25*, *PUB26*, *MYB6*, *VDAL*, *BIK1* or *GUS* was cloned into *pCAMBIA1300-ccLUC* and *pCAMBIA1300-GZnLUC*, respectively, and fused to *ccLUC* and *GZnLUC*, respectively. For *BD-PUB25*, *BD-PUB26*, *AD-MYB6s*, A*D-VDAL* or A*D-BIK1*, the CDS of *PUB25*, *PUB26*, *MYB6*, *VDAL* or *BIK1* was cloned into pGBKT7 or pGADT7, respectively. For *GST-PUB25*, *GST-PUB26*, *HIS-PUB26*, *GST-MYB6*, *MBP-MYB6*, *GST-VDAL*, *HIS-UBA1(E1)* and *HIS-UBC10(E2)*, the CDS of *PUB25*, *PUB26*, *MYB6*, *VDAL*, *E1* or *E2* was cloned into pGEX-4T-1, pET30a or pMAL-c5X, respectively, and fused to GST, HIS or MBP tag, respectively. For *BD-PUB25-pMet-VDAL*, *BD-PUB25-pMet-MYB6*, *BD-PUB26-pMet-VDAL* and *BD-PUB26-pMet-MYB6*, the CDS of *PUB25*, *PUB26*, *MYB6* or *VDAL* was cloned into MCSI or MCSII of pBridge vector, respectively.

### Phylogenetic Analysis

Protein sequence were obtained from National Center for Biotechnology Information (http://www.ncbi.nlm.nih.gov/). Full-length protein sequences were aligned and produced the phylogenetic tree by the neighbor-joining method of MEGA7.0.

### Pathogen Culture and Infection Assay

*V. dahliae* strain Vd991 was grown on PDA (potato dextrose agar) medium at room temperature (25 °C) for 1-2 weeks. The spore suspension solution was prepared at 1×10^6^ conidia/ml. Two-week-old *Arabidopsis* seedlings were removed carefully from pots and washed with water. The roots were dipped into the spore suspension solution for 5 min, and then re-planted in new prepared soil, and cultured at 23 °C in a 12-h-light/12-h-dark cycle for phenotyping. Disease index and pictures were recorded or taken on 10 −20 dpi (Fradin et al., 2011).

### Disease Index of Verticillium Wilt Caused by *V. dahlia*e

The disease index of *Arabidopsis* caused by *V. dahlia*e was recorded on 10-20 dpi. Since one of the distinguishing phenotypes after infected by *V. dahlia*e was the leaves turned to yellow abnormally and rapidly. According to the severity of the condition (proportion of leaves turned to yellow), disease was divided into 5 levels. The plant with no disease after incubation was defined as the level 0; then with less than 1/4 of leaves infected was defined as the level 1; with less than 1/2 of leaves infected was defined as the level 2; with less than 3/4 of leaves infected was defined as the level 3; and with more than 3/4 of leaves was defined as the level 4. 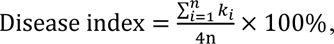 × 100%, k means the disease level of *Arabidopsis*; n means the number of *Arabidopsis*. Data were presented as means± SD, and analyzed by Student’s t test performed with Microsoft Excel and SPSS software. At least three biological replicates were used to perform each of the experiments, each replicate with at least 15 plants, and all the experiments were repeated for at least three times and the results were consistent.

### Confirmation of the Secreted Signal Peptide of VDAL in Yeasts

Functional validation of the VDAL signal peptide was performed using the yeast invertase secretion assay as described previously with slight modification (Oh et al., 2009). Briefly, the different form of VDAL signal peptide was introduced into pSUC2T7M13ORI (pSUC2-MSP) vector using *Eco*R1 and *Xho*1 restriction sites (pSUC2-VDAL). Yeast YTK12 strains were transformed with 500 ng pSUC2-VDAL or pSUC2-MSP using the lithium acetate method (Modrof et al., 2003), respectively. Transformed yeast cells were plated on selection CMD-W medium (0.67% yeast N base without amino acids, Trp dropout supplement, 2% sucrose, 0.1% glucose, and 2% agar). Positive clones were transferred to new CMD-W medium and YPRAA medium (1% yeast extract, 2% peptone, 2% raffinose, 2% agar, pH 5.8) for testing invertase secretion. Invertase enzymatic activity was assayed by 2,3,5-Triphenyltetrazolium chloride (Schenke et al., 2011) reduction assay. Untransformed YTK12 strain, YTK12 carrying the pSUC2-MSP and pSUC2-VDAL-SP vector were cultured in 5 mL YPDA liquid medium (2% peptone,1% yeast extract, 2% glucose, pH5.8) over night at 30 °C. Total yeast cells were collected and washed by ddH_2_O. Then the pellet was re-suspended with 0.1% colorless dye TTC at 30 °C for 1-2 h and color change was obtained in room temperature.

### Identification of VDAL-Interacting Proteins

To identify VDAL targeted protein in *Arabidopsis*, VDAL protein complex was purified as described previously with slight modification (Li et al., 2014). VDAL-MYC was expressed in *Arabidopsis* protoplasts. Total proteins were extracted in 8 mL Protein Extraction Buffer with 1% protease inhibitor (CWBIO). Cell debris was removed from the lysate by centrifugation at 12,000 g for 10 min. The supernatant was incubated with 150 μl anti-MYC agarose beads (sigma) for 2.5 h. Then the immunocomplexes were collected and washed with 1 mL low salt wash buffer (10 mM HEPES [pH 7.5], 100 mM NaCl, EDTA 1 mM, Glycerol 10%, 0.1% NP-40, 1 mM DTT, 1 mM PMSF and protease inhibitor cocktail) and high salt wash buffer (200 mM NaCl), each for four times. Immunocomplexes were eluted with Tris-glycine elution buffer. The total volume was concentrated to 30 μl with 3 kDa microspin columns. The 25 μl eluted proteins were loaded onto a single lane on a 10% SDS-PAGE gel for LC-MS (Beijing Protein Inovation) assay, another 5 μl eluted proteins were used for VDAL-MYC western blotting.

### Co-Immunoprecipitation

The protoplasts were transfected with the purified plasmids and incubated overnight (16 h). Total proteins were extracted for Co-IP with the extraction buffer (50 mM HEPES [pH 7.5], 150 mM NaCl, 1 mM EDTA, 0.5% Triton-X 100, 1 mM DTT, proteinase inhibitor cocktail). For anti-GFP IP, total proteins were incubated with 10 μl agarose conjugated anti-GFP antibodies (Chromotek) for 3 h and washed 3 times with the washing buffer (50 mM HEPES [pH 7.5], 150 mM NaCl, 1 mM EDTA, 0.5% Triton-X 100, 1 mM DTT). The simples were boiled for 5 min with 6×SDS loading buffer. The proteins were separated by SDS-PAGE and detected by anti-GFP, anti-MYC antibodies.

### Firefly Luciferase Complementation Imaging Assay

The full-length CDS sequence was fused to the C-terminus of GZnLUC and C-terminus of ccLUC (pCAMBIA1300). The constructed plasmids were transformed into Agrobacterium strain GV3101. Positive clones were incubated in 5 mL YEB liquid medium at 28°C for 16 h. The clones with different vectors were mixed equally (based on OD600). Agrobacteria were collected and resuspended in 2 mL of activity buffer (10 mM MES [pH 5.7], 10 mM MgCl_2_, 150 μM acetosyringone). After 2-5 h incubation at room temperature in darkness, the active Agrobacteria were infiltrated into expanded leaves of *N. benthamiana* and grown in the growth room for 48-72 h before the LUC activity measurement. For the CCD imaging and LUC activity measurement, infiltrated leaves were sprayed with 1 mM luciferin and then kept in the dark for 5 min. A pre-cold CCD (1300B, Roper) camera was used to capture the LUC image and the exposure time for signal collection was 15 min.

### Yeast Two-Hybrid Assay

To confirm the interaction between VDAL and PUB25 and PUB26, and the interaction between MYB6 and PUB25 and PUB26, full-length *PUB25*, *PUB26*, *MYB6* or *VDAL* CDS was fused into pGBKT7 (Fradin et al., 2011) and pGADT7 (AD) vectors, respectively. These plasmids were co-transformed into yeast strain Gold using the lithium acetate method (Kong et al., 2015). Transformed yeast cells were separately plated on 2D synthetic dropout medium (Trp-/Leu-) selective medium and incubated at 28 °C for 3-4 days. Then transformed yeast cells were transferred to fresh 2D and 4D media (Trp-/Leu-/His-/Ade-) with 1X, 10X, 100X and 1000X dilution for interaction detecting. The interaction of BIK1 and PUB26 was used as the positive control.

### The *in vitro* Ubiquitination Assay

GST-PUB25, GST-PUB26, GST-VDAL, MBP-MYB6, HIS-PUB26, HIS-UBA1 (E1) or HIS-UBC10 (E2) was expressed in *E. coli* strain BL21 or Rosetta, respectively. GST-PUB25, GST-PUB26 or GST-VDAL protein was purified on Glutathione Sepharose (GE), respectively. MBP-MYB6 protein was purified on amylose resin (NEB) and HIS-PUB26, HIS-UBA1 (E1) or HIS-UBC10 (E2) protein was purified on Ni Sepharose (GE), respectively.

For testing E3 ligase activity, 250 ng HIS-E1, 500 ng purified HIS-E2, 1.25 µg Flag tagged ubiquitin (Boston Biochem), 2 µg purified GST-PUB25 and PUB26 or HIS-PUB26 were added to 30 µL of ubiquitination reaction buffer (50 mM Tris-HCl pH 7.5, 2 mM ATP, 5 mM MgCl_2_, 2 mM DTT). After 3 h at 30 °C, all reactions were stopped by adding 6×SDS loading buffer, and boiled for 5 min.

For GST-VDAL and MBP-MYB6 ubiquitination assay, 1 µg purified GST-VDAL or MBP-MYB6 was added to above 30 µL of ubiquitination reaction buffer. After incubated for 3 h at 30 °C, all reactions were stopped by adding 6×SDS loading buffer, and boiled for 5 min. For competitive ubiquitination assay, 1 µg purified MBP-MYB6 and different concentration (1 µg, 3 µg, 7 µg) of GST-VDAL, or 1 µg purified GST-VDAL and different concentration (1 µg, 3 µg, 7 µg) of MBP-MYB6 were added to above 30 µL ubiquitination reaction buffer. After incubated for 3 h at 30 °C, all reactions were stopped by adding 6×SDS loading buffer, and boiled for 5 min. The proteins were separated by 10% SDS-PAGE gel, and detected with anti-Ub, anti-GST or anti-MBP antibodies.

### The *in vivo* Ubiquitination Assay

To determine the specificity for protein degradation of VDAL, *in vivo* ubiquitination assay was performed. Total proteins of 10 day VDAL-MYC in Col and *pub25 pub26* double mutant were extracted with the extraction buffer (50 mM HEPES [pH 7.5], 150 mM NaCl, 1 mM EDTA, 0.5% Triton-X 100, 1 mM DTT, proteinase inhibitor cocktail), and incubated with 20 μl MYC beads coated with MYC antibodies (Sigma, A7470) for 3 hrs, and washed three times with washing buffer (50 mM HEPES [pH 7.5], 150 mM NaCl, 1 mM EDTA, 0.5% Triton-X 100, 1 mM DTT). The simples were boiled for 5 min with 6×SDS loading buffer. The proteins were separated by SDS-PAGE, and detected by anti-Ub and anti-MYC antibodies.

### Protein Pull-Down Assay

To confirm the interaction between PUBs and VDAL, as well as PUBs and MYB6, the pull-down assay was performed as described previously (Wang et al., 2018b) with slight modifications. Briefly, GST, or GST-PUB26 protein (10 µg) was incubated with Glutathione Sepharose for 2 h with 1 mL pull-down binding buffer (150 mM NaCl, 10 mM Na_2_HPO_4_, 2 mM KH_2_PO_4_, 2.7 mM KCl, and 0.1% Nonidet P-40, pH 7.4) at 4 °C. After a centrifugation at 1000 g for 3 min at 4 °C, the buffer was removed and 1 mL fresh binding buffer was added to the Glutathione Sepharose, then 2 µg VDAL-HIS, or MBP-MYB6 protein was added to the above fresh binding buffer (then 2 µg of VDAL-HIS together with 2 µg MBP-MYB6 protein was added to the above fresh binding buffer), along with Sepharose. The tube was rotated at 4 °C for 2 h for protein binding. After a centrifugation at 1000 g for 3 min at 4 °C, the buffer was removed, and the Sepharose was washed five times with 1× PBS (150 mM NaCl, 10 mM Na_2_HPO_4_, 2 mM KH_2_PO_4_, and 2.7 mM KCl, pH 7.4) to remove non-specifically bound proteins. All reactions were stopped by adding 50 mL 1× PBS and 10 µl 5X SDS loading buffer boiled for 5 min. The proteins were separated by 10% SDS-PAGE gel and detected with anti-Ub, anti-GST or anti-MBP antibodies. All reactions were stopped by adding 50 μl 1× PBS and 10 µl 5X SDS loading buffer, and boiled for 5 min. The proteins were separated by 10% SDS-PAGE gel, and detected with anti-HIS, anti-GST or anti-MBP antibodies.

### ROS Burst Assay

ROS burst assay was performed as described previously with slight modifications (Wang et al., 2018a). Briefly, 5 mm-diameter leaf disces were collected from 4-5 week-old plants into 96-well plates, and floated overnight on sterile water in a 96-well plate. In the following day the water was replaced with a solution of 20 mM luminol (Sigma) and 10 mg/mL horseradish peroxidase (Sigma) containing different concentration of VDAL, then luminescence recorded by the Tecan-i-control outfitted with the LUM module.

### Cell-Free Protein Degradation Assay

To identify the protein stability of VDAL and MYB6, the cell-free protein degradation assay was performed as described previously (Kong et al., 2015) with slight modifications. For protein degradation in transgenic plants, the seeds of *Pro35S::VDAL-GFP* grown at 22°C for 10 days under long-day condition (16 h light/8 h dark cycles), then total proteins were extracted with native protein extraction buffer [50 mM Tris-MES (pH 8.0), 0.5 M sucrose, 1 mM MgCl2, 10 mM EDTA (pH 8.0),5 mM DTT], the extracted supernatants were divided into two or four equal parts with addition of 50 µM MG132, 1 mM ATP or not, respectively, and the samples were cultured at 22°C with shaking for different times. For purified protein degradation, the seeds of Col and *pub25* and *pub26* grown at 22 °C for 10 days under long-day condition (16 h light/8 h dark cycles), then total proteins were extracted with native protein extraction buffer, 200 ng purified protein of GST-VDAL or GST-MYB6 from *E. coli* strain Rosetta was incubated in 600 μg total proteins for each reaction with addition of 50 µM MG132 and 1 mM ATP, or 1 mM ATP only, then the reactions were cultured at 22 °C for different times. All reactions were stopped by adding 5×SDS loading buffer with boiled for 5 min. The proteins were separated by 10% SDS-PAGE gel and detected with anti-GFP or anti-GST antibodies.

### Chitin Induced MAPK Kinase Activities Assay

MAPK kinase activities assay was performed as described previously with slight modification (Yamada et al., 2016; Wang et al., 2017a). Briefly, *Arabidopsis* seedlings were grown on MS medium for 7 days under 22 h light/2 h dark at 22 °C and then treated with or without 500μg/mL chitin 10 minutes. Total proteins were extracted with the extraction buffer (50 mM HEPES [pH 7.5], 150 mM NaCl, 1 mM EDTA, 0.5% Triton-X 100, 1 mM DTT, proteinase inhibitor cocktail, 1× EDTA-free protease inhibitor cocktail, and 1× Phostop phosphatase inhibitor cocktail). The proteins were separated by SDS-PAGE and detected by anti-phospho-p44/42 MAPK antibodies (Cell Signaling #4370) and anti-actin antibodies respectively.

### Subcellular Localization and GUS Staining

To identify the subcellular localization, the full-length CDS of *VDAL* was cloned into PCAMBIA1300 vector under control of *35S* promoter and fused to GFP tag. To determine the subcellular localization of VDA in cells, the protoplasts were transfected with each purified plasmid and incubated overnight (16 h). The GFP images were acquired at excitation of 488 nm and emission of 525 nm. To determine the subcellular localization of VDAL in the transgenic plants, the *Pro35S::VDAL-GFP* homozygous transgenic seedlings were obtained via the floral dip method (Clough and Bent, 1998), The GFP images were acquired at excitation of 488 nm and emission of 525 nm.

To identify the expression of *PUB2*5, *PUB26* and *MYB6* in plants, the promoter of *PUB2*5, *PUB26* and *MYB6* was cloned into pCAMBIA1391 vector respectively. The plasmid was subsequently transformed into Agrobacterium GV3101 and transferred into plants via the floral dip method. T3 homozygous transgenic seedlings were used for the GUS staining assay.

### VDAL Zinc-Binding Assay

To determine the zinc-binding activity of VDAL, zinc-binding assay was performed as described previously (Citiulo et al., 2012) with slight modifications. 4 ng GST or GST-VDAL purified protein were loaded onto 10 kDa microspin columns (Ambion). After washed twice with 1 mL HS buffer (50 mM HEPES-KOH pH 7.5, 200 mM NaCl), supernatants were transferred to reaction tubes and incubated with 0.1 mM Zn^2+^ for 1 h at 37 °C. The zinc-loaded proteins were then transferred to 10 kDa microspin columns and sequentially washed with 4 mL of HS buffer. Each flow-through was assayed for zinc content by PAR assay, until there no Zn^2+^ in the buffer. Briefly, 4-(2-pyridylazo) resorcinol (PAR) was added to each sample to 0.1 mM and optical density measured at 490 nm against a Zn^2+^ standard curve. Following 15 washes, undigested and digested (40 mg proteinase K, 50 °C for 30 min) samples were again assayed for zinc content by PAR assay.

### Biacore Assay

To determine the affinity of HIS-VDAL to GST-PUB26, or MBP-MYB6 to GST-PUB26, Biacore assay was employed. Immobilization of anti-HIS & anti-GST antibodies onto different CM5 chips: Anti-GST was immobilized respectively at flow cell 1, 2, and the immobilization level of each flow cell was 4500 Ru, 5500 Ru. Anti-HIS was immobilized respectively at flow cell 1, 3, and the immobilization level of each flow cell was 10000 Ru. PBS + 0.05% [v/v] surfactant P20 was used as the running buffer.

To test the affinity of HIS-VDAL to GST-PUB6, HIS-VDAL-DSP captured on flow cell 3 of anti-HIS chip. Analyses GST-PUB26, MBP-MYB6, GST, MBP flowed through flow cell 1, 2, 3, 4 at the Flow rate 10 μL/min and flow cell 1 as the control. Association time:120 s for GST-PUB26, MBP-MYB6; Disassociation time: 300 s for MBP-MYB6; 60 s for GST-PUB26; Analysis concentration: MBP-MYB6 (μg/mL): 0, 3.25, 7.5, 15,.30, 60; GST-PUB26 (μg/mL): 0, 2.18, 4.375, 8.75, 17.5, 35. 2.

To determine the affinity of GST-PUB26 to MBP-MYB6, GST-PUB26 was captured on flow cell 2 of anti-GST chip. Analyses MBP-MYB6, MBP flowed through flow cell 1, 2 at the Flow rate 10 μL/min and flow cell 1 as the control. Association time: 120 s for MBP-MYB6; Disassociation time: 300s for MBP-MYB6; Analysis concentration: MBP-MYB6 (μg/mL): 0, 7.5, 15,.30, 60, 120. Regeneration: Glycine-HCl pH 1.5 was as regeneration at 30 μL/min about 60s. Biacore T200 evaluation 3.1 was used for data analysis. The generated signals were subtracted from reference channel and then analyzed with kinetics fitting.

### Quantitative RT-PCR

14 day-old seedlings were treated with or without 1×10^6^ conidia/mL spore suspensions of *V. dahliae* 12 h. Total RNAs were extracted by HiPure Plant RNA Mini Kit RNeasy Plant Mini Kit (Magen,cat.R4151) with the kit instructions. The RNAs were reversely transcribed to cDNAs with MMLV reverse transcriptase (Thermo). Quantitative RT-PCR was performed with a 7300 Real-Time PCR system (Applied Biosystems) using SYBR Premix Ex Taq (TaKaRa). *Actin* was employed as an internal control.

### RNA-Seq Analysis

*Arabidopsis* seedlings were grown on MS medium for 10 days under 22 h light/2 h dark at 22 °C and then treated with or without 1×10^6^ conidia/mL spore suspensions of *V. dahliae* for 12 h. Total RNAs were extracted by HiPure Plant RNA Mini Kit RNeasy Plant Mini Kit (Magen, cat. R4151) with the instruction. 3 μg RNAs for each sample were used for library construction using NEB Next Ultra RNA Library Prep Kit for Illumina (NEB, USA, cat. no. E7530) according to the instruction and RNA-Seq on Illumina Hiseq 2500 platform.

About 3.0 GB clean reads were generated for each sample. All reads were trimmed to 150-bp paired-end reads with high quality according to the base quality. The trimmed reads were mapped to the A. thaliana reference genome (TAIR10) using bowtie2 with default settings. Differential gene expression analysis was performed using edgeR (Robinson et al., 2010).

Fold changes of the genes induced significantly by *V. dahliae* treatment were compared with the control sample in each group (using P<0.05, t-test and |logFC| >1). Genes significantly induced by *V. dahliae* in the Col group were chosen for the comparison with expression levels of the treatment samples between different groups. To compare the change levels of the treatment samples between different groups, we calculated relative expression level (using P<0.05 or *P*<0.01,t-test). Fold change of each gene in the treatment samples in *myb6-Cas9*, *pub25 pub26* or *VDAL-OE* minus the fold change of the same gene in the treatment samples in Col was considered as the relative expression level of the gene in *myb6-Cas9*, *pub25 pub26* or *VDAL-OE* comparing to Col. Thus positive or negative value of the relative expression level indicated the change level of the gene in *myb6-Cas9*, *pub25 pub26* or *VDAL-OE* was higher or lower than the change level in Col. Then Venn diagrams showing the differentially regulated genes (down regulated genes and up regulated genes) in common between the different sample were generated using an online tool jvenn (http://jvenn.toulouse.inra.fr/app/example.html). The heat map was drawn according to the expression levels in the *myb6-Cas9*, *VDAL-OE* and *PUB25* and *PUB26* compared to Col using in-house python scripts. Gene enrichment analysis were generated by online tool AGRI GO (http://bioinfo.cau.edu.cn/agriGO/index.php).

Our raw data and the processed RNA-seq data have been deposited in the National Center for Biotechnology Information Gene Expression Omnibus (https://www.ncbi.nlm.nih.gov/sra/PRJNA636958).

### Virus-Induced Gene Silencing (VIGS)

Agrobacterium cultures containing pTRV-CLA1, pTRV-GhMYB330 and pTRV-GhPUB26, pTRV-GFP and pTRV-RNA1 were cultured in 2-3 mL LB at 37 °C 12-16 h, then transferred to the 100 mL LB continue to cultured 12-16 h, cultures were collected and resuspended to OD600=0.8 with infiltration buffer (10 mM MES, 10 mM MgCl_2_, 200 μM acetosyringone, and pH5.4), set in dark 3 h. GV3101 cells containing pTRV-RNA1 were mixed with cells harboring pTRV-GhMYB330, pTRV-GhPUB26 and pTRV-GFP and infiltrated into the true leaves ofcotton seedlings. After two weeks, the 2 mL spore suspension solution of *V. dahliae* was prepared at 1×10^6^ conidia/mL were infiltrated into the stem near the cotyledon. Disease index and pictures were recorded or taken on 10 d-25 d after incubation.

### Accession Numbers

Sequence data from this article can be found in the GenBank/EMBL data libraries under the following accession numbers: *PUB25*, *AT3G19380*; *PUB26*, *AT1G49780*; *MYB6*, *AT4G09460*; *VDAL*, *AY524791*, *GhPUB26*, *GH_D13G0403*; *GhMYB330*, *GH_A13G1430*. All transgenic *Arabidopsis thaliana* lines and T-DNA insertion mutants used in this study are in the Columbia-0 (Col-0) ecotype background. The T-DNA insertion mutants *pub25* (SALK-147032C), *pub26* (S351943,GK308D07), *myb6-1* (SALK_074789C) and *myb6-2* (CS403018) were obtained from the ABRC. *myb6*-CRISPR Cas9 mutant was obtained though CRISPR Cas9 system.

## Supplemental Data

Supplemental Figure 1. Sequence analysis of VDAL.

Supplemental Figure 2. Phylogenetic tree of the VDAL from different fungi

Supplemental Figure 3. PUB25/26 interact with different VDAL fragment in yeast two-hybrid system.

Supplemental Figure 4. PUB25 and PUB26 have auto-ubiquitination activity and PUB26 can ubiquitinate VDAL *in vitro*

Supplemental Figure 5.*pub25 pub26* double mutant shows resistant to Verticillium wilt caused by *V. dahliae*.

Supplemental Figure 6. MYB6 is stable in *pub25 pub26* double mutant.

Supplemental Figure 7. VDAL and MYB6 are competitors in PUB25 ubiquitination

Supplemental Figure 8. The gene expression analyses in VDAL transgenic plants and different mutants

Supplemental Table 1 Affinity and kinetics parameters of the target proteins from Biacore assay.

Supplemental Data Set 1 VDAL interaction protein candidates by LC-MS.

Supplemental Data Set 2 PUB26-ARM yeast two-hybrid screen candidates.

Supplemental Data Set 3 Gene expression analysis from RNA-seq.

Supplemental Data Set 4 overlapped genes in *myb6* down-regulated and *pub25 pub26* and *VDAL-OE* up-regulated genes.

Supplemental Data Set 5 overlapped genes in *myb6* up-regulated and *pub25/26* and *VDAL-OE* down-regulated genes.

Supplemental Data Set 6 Primers used in this research.

Supplemental data 1 The sequence and Accession numbers of proteins used to Phylogenetic tree analysis.

## ACKNOWLEDGMENTS

This research was financially supported by the National Key Research and Development Program of China (grant 2016YFD0101904); Major transgenic project Program of China (2016ZX08005-004, 2014ZX08005-02B).

We thank Prof. Dingzhong Tang at Fujian Agriculture and Forestry University, Prof. Xianzhong Feng, Dr. Jiantian Leng and Yu Hui, at Northeast Geography and Agroecology for providing valuable advices on writing of manuscript; Prof. Ruyu Li and Yongqing Li at South China Botanical Garden, Chinese Academy of Sciences for donating cotton cultivar; Prof. Zhaosheng Kong and Dr. Zhidi Feng at Institute of Microbiology, Chinese Academy of Sciences for donating Verticillium dahliae strain. Libo Shan at Texas A&M University for donating pSUC2 vector. We thank Prof. Jianmin Zhou at State Key Laboratory of Plant Genomics, China Institute of Genetics and Developmental Biology, Chinese Academy of Sciences for providing *NP-PUB25-Flag* seeds and valuable advice.

## AUTHOR CONTRIBUTIONS

Aifang Ma, Z.G.,, and J.Q. designed the study. A. M. and D.Z. performed the experiments. A. M., H.L., Y.G., Y.H., Z.G., and J.Q. analyzed the data. A. M., Z.G. and J.Q. wrote the paper. All other authors participated in the discussion of the results and commented on the manuscript.

